# Distinct In Vivo Electrophysiological Profiles of Mediodorsal Thalamus Subdivisions

**DOI:** 10.64898/2026.09.02.748524

**Authors:** Katerina Panti, Charlotte J. Stagg, Andrew Sharott, Jeffrey Stedehouder

## Abstract

The mediodorsal nucleus of the thalamus (MD) plays a key role in complex cognitive processes, and its dysfunction is linked to various neurological and psychiatric conditions. The MD is divided into three parts that vary in anatomical connectivity, molecular expression, and *ex vivo* electrophysiology. Here, we describe *in vivo* single-neuron electrophysiological recordings across the three MD subdivisions in head-fixed, behaving, adult wildtype mice. We report large differences in extracellular waveforms, spiking activity, and burst firing characteristics across MD subdivisions. Specifically, central MD subdivision (MDc) neurons showed markedly increased waveform amplitude, higher spontaneous firing rate, and increased burst firing compared to medial (MDm) and lateral (MDl) subdivision neurons. Single-neuron electrophysiology features were sufficient to classify neurons into respective anatomical subdivisions. Hierarchical clustering revealed MDc is distinct from MDl and MDm and more akin to other, non-MD thalamic nuclei. Together, these data suggest distinct *in vivo* electrophysiological profiles of MD subdivisions, posing implications for investigating MD thalamus in health and disease.

**HIGHLIGHTS:**

- Single-neuron recordings across subdivisions of mediodorsal thalamus in awake mice
- Large differences in waveform, spiking, and burst activity metrics across the three subdivisions
- Single-neuron electrophysiological profiles sufficient to predict subdivision location
- The central subdivision is largely distinct from the medial and lateral subdivisions

## INTRODUCTION

The mediodorsal nucleus of the thalamus (MD) is the second largest thalamic nucleus of the human brain and has classically been considered part of higher-order thalamus.^1, 2^ The MD has been identified as a critical region for various cognitive processes including working memory^3, 4^, attention^3, 5, 6^, and goal-directed behavior^7, 8^, as well as emotional processing^9, 10^. Converging clinical and preclinical evidence implicate the MD in psychiatric and neurological conditions, such as schizophrenia^11, 12^, depression^13, 14^, fatal familial insomnia^15^, epilepsy^16–18^, autism spectrum disorder^19^, chronic pain^20^, Alzheimer’s disease^21^, and Parkinson’s disease^22^. Notably, modulation of the MD is increasingly recognized as a potential therapeutic target for schizophrenia^23, 24^, epilepsy^25^, and apathy in Parkinson’s disease^26^.

The MD is a heterogeneous structure that increases in size with phylogenetic evolution, being most developed in primates, especially in humans^1^. In rodents, the MD is commonly divided into three parts across its mediolateral axis including a medial part (MDm), a central part (MDc), and a lateral part (MDl)^1, 27, 28^. Across species, the MD includes further subgroupings within these three main subdivisions^29, 30^, but overall the MD is broadly homologous across rodents and primates^1^. Converging lines of evidence have suggested variation across MD subdivisions at the level of anatomical inputoutput connectivity, molecular expression, and *ex vivo* electrophysiology^31–35^. Anatomically, MD subdivisions have dense connections to specific sub-regions that span the entire frontal cortex and parts of temporal cortex and receive distinct modulatory subcortical inputs^3, 27, 28, 32, 36–40^. MDm primarily receives inputs from amygdala, ventral pallidum, and nucleus accumbens and reciprocally connects to ventromedial PFC^3, 27, 28, 32, 36, 37, 39, 40^. MDc receives inputs from the globus pallidus and endopiriform nucleus and forms a circuit with dorsolateral PFC and orbitofrontal cortex (OFC)^3, 27, 28, 32, 36–40^. MDl receives inputs from the substantia nigra pars reticulata and brainstem and reciprocally connects to dorsomedial PFC and anterior cingulate cortex (ACC)^3, 27, 28, 32, 36, 37, 40^. Furthermore, MD subdivisions have been shown to contain different gene expression profiles^34, 41, 42^ while electrophysiological differences in membrane properties of single neurons measured with *ex vivo* intracellular recordings have also been reported^31, 33, 35^. Functionally, studies in non-human primates^43, 44^ and rodents^3, 5, 33^ have suggested that MD subdivisions show dissociable roles in supporting frontal cortical networks during specific cognitive tasks. Together, these findings suggest that the three MD subdivisions present different characteristics across multiple anatomical, molecular, and *ex vivo* electrophysiological features but whether these translate to divergent *in vivo* electrophysiological properties remains unknown.

Here, we describe high-density *in vivo* electrophysiological recordings of thalamus across the three MD subdivisions in head-fixed, behaving, wildtype male mice. We report large differences between MDm, MDc, and MDl in extracellular waveform, spiking, and burst firing metrics across independent cohorts. *In vivo* electrophysiological measures were sufficient to classify neurons into MD subdivisions while hierarchical clustering indicated MDc as distinct from MDm and MDl. Hierarchical clustering including additional thalamic nuclei further revealed that MDc was more akin to other, non-MD nuclei. Thus, these data suggest that subdivisions of rodent MD thalamus show distinct *in vivo* electrophysiological profiles. These findings extend the parameter space of differences between the three MD subdivisions and allow insights to guide future investigations targeting the MD in health and for neurological and psychiatric disorders.

## RESULTS

### *In vivo* electrophysiological recordings across rodent mediodorsal thalamus subdivisions

To examine *in vivo* single-neuron electrophysiological characteristics across MD subdivisions, we performed large-scale neural recordings with Neuropixels 1.0 probes across 14 adult, male wildtype C57BL/6J mice (**Fig. 1a**; **Supplementary Fig. 1a**; **Methods**). We focused our analysis on recording sessions that included MD thalamus (3 cohorts, 12 animals, 34 recording sessions, **Methods**) (**Fig. 1b**; **Supplementary Fig. 1b-d**). Data for one cohort (‘Cohort 2’) including MD thalamus have been previously described elsewhere^12^ and were re-analyzed here. Across all cohorts, recordings were obtained acutely from head-fixed behaving mice performing a sensorimotor task over several days (**Supplementary Fig. 1e-h**). Cohorts had minor differences in behavioral task parameters and performance, craniotomy location, and probe insertion location, angle, and depth (**Methods**; **Supplementary Fig. 1**). Probes targeted to the MD were acutely inserted downward into the brain and medially directed inwards (∼5-10°) over the right hemisphere through cortex, hippocampus, and habenula to reach thalamus (**Fig. 1c**). We employed histological reconstruction using fluorescent DiI dye on the probes as well as a combination of previously validated electrophysiological landmarks (e.g. high unit density in CA1 pyramidal cell layer, no units in transition border from hippocampus to thalamus) to align probe channels linearly with distinct brain areas (**Fig. 1d**; **Methods**). MD thalamus was either the deepest recorded brain region (∼40% of insertions; 14 out of 34 probe insertions) or probes passed through the MD with the probe tip residing in more ventral thalamus or regions of the hypothalamus (∼60% of insertions; 20 out of 34 probe insertions).

**Figure 1.**
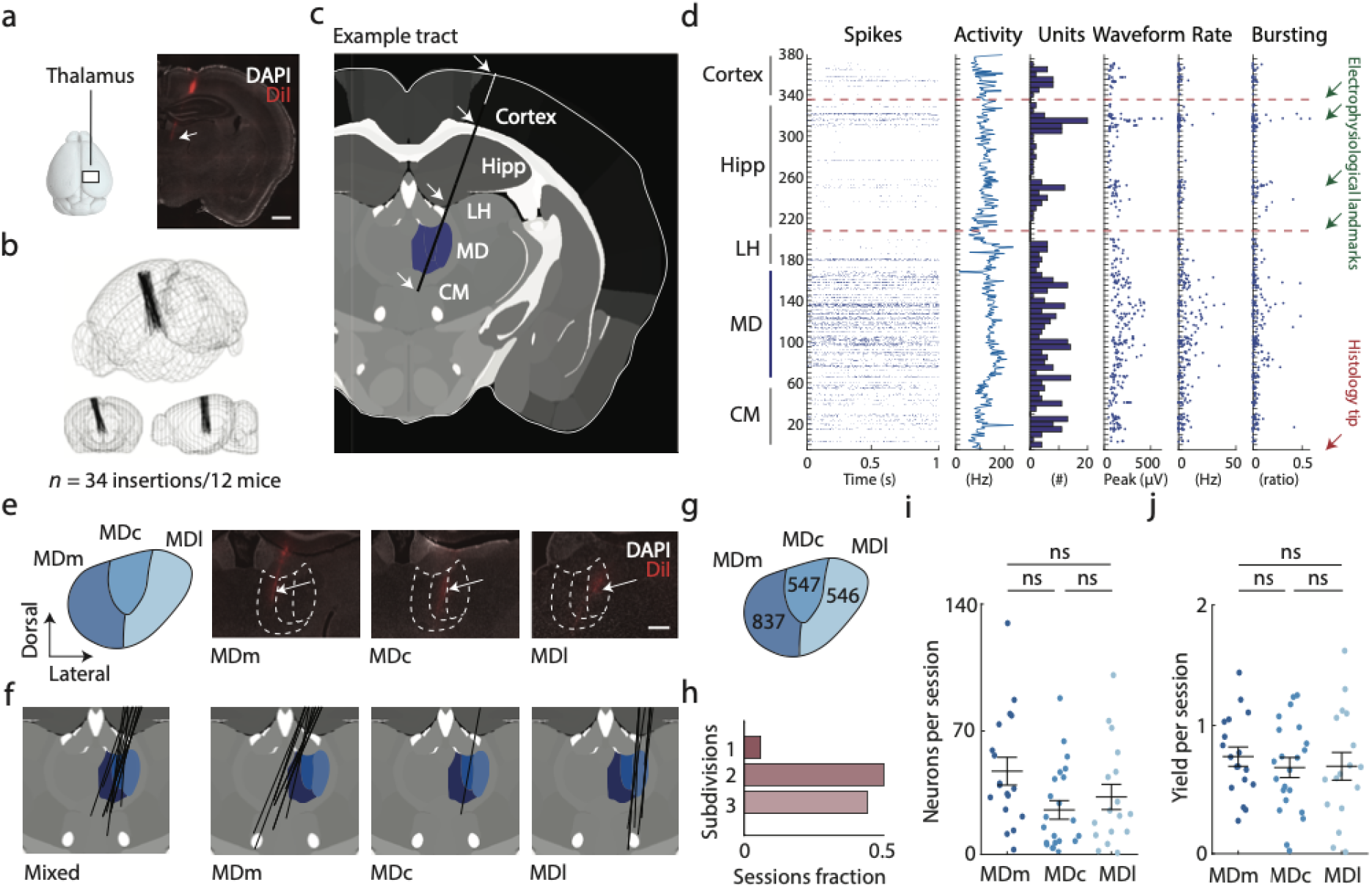
*In vivo* electrophysiological recordings of rodent mediodorsal thalamus subdivisions. **(a)** *Left*: *In vivo* electrophysiological recordings in C57BL/6J mice using Neuropixels probes targeted to thalamus. *Right*: Exemplar low-magnification coronal epifluorescent microscope image of probe track (arrow) passing through the mediodorsal (MD) nucleus of thalamus. DAPI nuclear marker in white, DiI probe track in red. Scale bar, 1 mm. **(b)** 3D schematic of brains showing all reconstructed probe trajectories through MD. Trajectories across three independent cohorts in **Supplementary Fig. 1**. **(c)** Example coronal view of an insertion track of a Neuropixels probe through MD thalamus with traversed brain regions indicated in text and landmarks indicated with arrows. MD thalamus highlighted in blue. **(d)** Example region alignment for the single probe insertion shown in (c). Information on spiking, waveform, and bursting activity across channel depth (top to bottom) through cortex, hippocampus, lateral habenula, MD thalamus, and centromedian thalamus. Red lines indicate landmarks for linearly rearranging channels for brain region borders. Green arrows represent previously validated electrophysiological landmarks used to rearrange channels (see **Methods**). **(e)** *Left:* Schematic representation of MD nucleus segregated into subdivisions (medial MDm, central MDc, and lateral MDl) in the right hemisphere with different shades of blue. *Right:* Exemplar high-magnification coronal epifluorescent microscope images of probe tracks (arrows) in central, medial and lateral subdivisions of the MD. DAPI nuclear marker in white, DiI probe track in red. Scale bar, 200 μm. **(f)** *Left*: Schematic coronal visualizations of probe insertions (black lines) that pass through multiple MD subdivisions simultaneously (*n* = 19 recording sessions). *Right*: Probe insertions that exclusively pass through only MDm (*n* = 8 sessions), MDc (*n* = 1 sessions), and MDl (*n* = 6 sessions), respectively. MD thalamus subdivisions highlighted in shades of blue. **(g)** Total number of recorded neurons for each MD subdivision following quality control and manual curation. Further details across the three cohorts in **Supplementary Fig.7**. **(h)** Fraction of sessions with single, two, and three simultaneously recorded MD subdivisions from a single Neuropixels probe recording. 50% of recordings covered two or three MD subdivisions. **(i)** Comparison of recorded number of neurons per session across MDm (*n* = 18), MDc (*n* = 21), and MDl (*n* = 16) shows no differences between subdivisions. **(j)** Comparison of yield (number of neurons/number of channels) per session across MDm (*n* = 18), MDc (*n* = 21), and MDl (*n* = 16) shows no differences between sessions. Black lines in (i) and (j) indicate means. Error bars denote standard error of the mean. n.s *P* > 0.05. One-way ANOVA followed by *post hoc* Tukey-Kramer comparisons in (i) and (j). Hipp hippocampus, LH lateral habenula, CM centromedian thalamus, MD mediodorsal thalamus. Statistics in **Supplementary Table 2**.

Following automated and manual curation of electrophysiological data (**Methods**), we retained a total of 1930 units from MD thalamus. Units (‘neurons’) were assigned into three subdivisions along the right mediolateral axis (MDm, MDc, and MDl) using a customized atlas (**Fig. 1e, f**; **Supplementary Fig. 2**; **Methods**). From these assignments, 837 neurons were located within MDm (∼43.4%), 547 neurons within MDc (∼28.3%), and 546 neurons within MDl (∼28.3%) (**Fig. 1g**; **Supplementary Table 1**). In 15 recording sessions (∼44%), neurons were recorded from a single subdivision, in 17 sessions (50%) neurons were recorded from two MD subunits simultaneously, and in two sessions (∼6%) neurons were recorded from all three subdivisions simultaneously with a single probe (**Fig. 1f, h**). No major differences were present across MD subdivisions in quality control processing (**Supplementary Fig. 3a-e**) or in the number of recorded neurons per session, recorded neurons per animal, yield per session, or yield per animal (**Fig. 1i, j**; **Supplementary Fig. 3f-g**). Overall, this experimental approach provided high-quality, single-neuron *in vivo* electrophysiological recordings across the three subdivisions of MD thalamus in head-fixed behaving mice.

### Mediodorsal thalamus subdivisions show large differences in waveform metrics

Examination of extracellular single-channel spike waveforms can indicate differences in the underlying morphoelectrical properties of individual neurons. Thus, we first examined waveform metrics of individual neurons across MD subdivisions obtained from non-filtered single-channel refined extracellular spike waveforms (**Fig. 2a, b**; **Methods**). Waveform amplitude was different between neurons across MD subdivisions (MDm: 160.2 ± 3.3 μV, MDc: 193.2 ± 4.0 μV, MDl: 159.8 ± 3.1 μV, χ^2^= 86.7, *P* < 0.001, Kruskal-Wallis test) with MDc neurons displaying larger amplitudes than both MDm and MDl neurons (+20.6%, *P* < 0.001, and +20.9%, *P* < 0.001, respectively, *post-hoc* Dunn’s test for multiple comparisons, **Fig. 2c**). In addition, trough-to-peak latency (repolarization latency) followed a unimodal distribution in all subdivisions and differed across MD subdivisions (MDm: 0.51 ± 0.01 ms, MDc: 0.48 ± 0.01 ms, MDl: 0.50 ± 0.01 ms, χ^2^= 83.1, *P* < 0.001, Kruskal-Wallis test, **Fig. 2d**), with MDc neurons displaying more narrow waveforms compared to MDm and MDl neurons (-6.9%, *P* < 0.001, and -3.4%, *P* < 0.001, respectively, *post-hoc* Dunn’s test for multiple comparisons, **Fig. 2d**). Additional wave-form differences were observed in peak-trough ratio (MDm: 0.46 ± 0.01, MDc: 0.49 ± 0.01, MDl: 0.48 ± 0.01, χ^2^= 50.4, *P* < 0.001, Kruskal-Wallis test), and shoulder asymmetry (MDm: 0.90 ± 0.01, MDc: 0.92 ± 0.01, MDl: 0.90 ± 0.01, χ^2^ = 11.5, *P* = 0.010, Kruskal-Wallis test, **Fig. 2e, f**). Subsequent analyses of high-pass filtered waveforms (**Supplementary Fig. 4a; Methods**) to exclude effects of slower oscillations, yielded similarly different results across MD subdivisions for all metrics except for shoulder asymmetry (**Supplementary Fig. 4b-e**).

**Figure 2.**
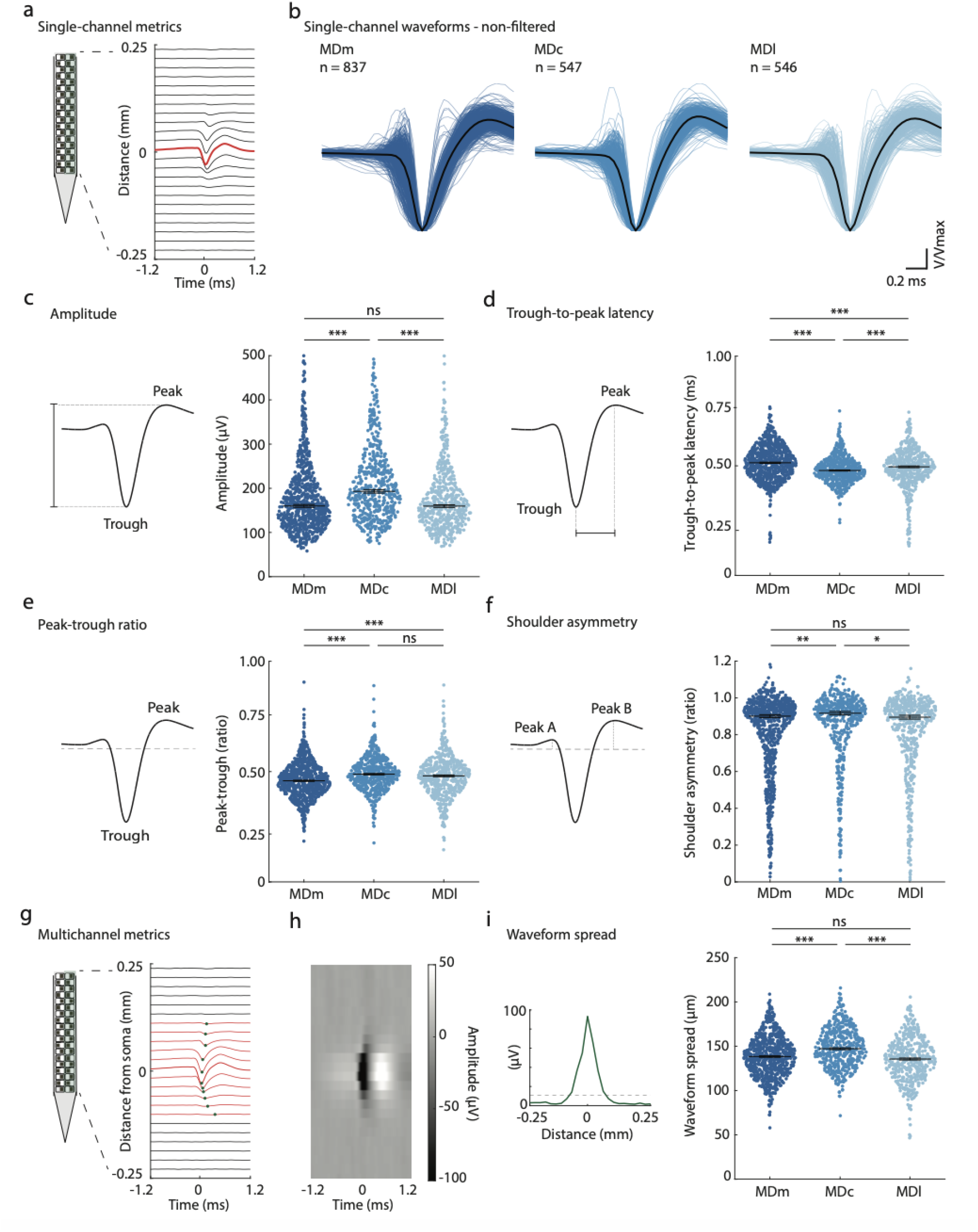
Mediodorsal thalamus subdivisions show differences in waveform metrics. **(a)** Illustration of single-channel extracellular waveform of an example neuron. For each neuron, we analyzed waveform metrics from best channel with maximum amplitude (in red; see **Methods**). **(b)** Non-filtered, single-channel refined waveforms of individual neurons in MDm (left), MDc (mid), and MDl (right). Black waveform indicates mean of all waveforms. **(c)** *Left:* Schematic of extracellular waveform amplitude calculation. *Right*: Swarm plot comparing amplitude across neurons MD subdivisions, showing differences between subdivisions. Y-axis capped at 500 µV for clarity. **(d)** *Left:* Schematic of trough-to-peak latency calculation. *Right*: Swarm plot comparing trough-to-peak latency across MD subdivisions, showing differences between subdivisions. **(e)** *Left:* Schematic of peak-trough ratio calculation. *Right*: Swarm plot comparing peak-trough ratio across MD subdivisions, showing differences between subdivisions. **(f)** *Left:* Schematic of shoulder asymmetry calculation. *Right*: Swarm plot comparing shoulder asymmetry across MD subdivisions, showing differences between subdivisions. **(g)** Schematic of multichannel extracellular waveform of an example neuron. For each neuron, we analyzed waveforms measures on one column of the probe (see **Methods**). The multichannel waveform includes the channel with the maximum amplitude and up to 12 channels both above and below (red lines). Green dots indicate waveform trough at each recording site. **(h)** Heatmap where each row represents the same waveform from one channel over time, with intensity reflecting waveform amplitude. **(i)** *Left*: Visualization of peak amplitude from (h), showing waveform spread across channels away from best channel. Dashed line indicates 12% from peak amplitude and represents the spread threshold. *Right*: Swarm plot comparing waveform spread across MD subdivisions, showing differences between subdivisions. Blue dots indicate individual neurons. Black lines in (c) through (f) indicate medians. Black line in (i) indicates mean. Error bars denote standard error of the mean. *** *P* < 0.001; ** *P* < 0.01; * *P* < 0.05; ns non-significant *P* > 0.05. In (c) through (f): Kruskal-Wallis test followed by Dunn’s multiple comparisons. In (i): One-way ANOVA test followed by *post hoc* Tukey-Kramer comparisons. V waveform extracellular voltage; Vmax waveform peak voltage. Statistics in **Supplementary Table 2.**

The use of Neuropixels probes with linearly arranged high-density channels allows for examination of multichannel waveform features, providing an additional characterization dimension of individual neurons^45^. Thus, we next examined multichannel features of extracellular spike waveforms across MD subdivisions (**Fig 2. g, h**; **Methods**). Waveform spread along the probe differed across MD subdivisions (MDm: 138.6 ± 0.8 μm, MDc: 148.8 ± 1.1 μm, MDl: 135.7 ± 1.1 μm, *F* (2, 1708) = 42.85, *P* < 0.001, One-way ANOVA), with MDc neurons showing larger waveform spread compared to both MDm and MDl neurons (+7.3% *P* < 0.001, and +9.6%, *P* < 0.001, respectively*, post-hoc* Tukey-Kramer test for multiple comparisons, **Fig. 2i**). Similar differential effects across MD subdivisions were observed for the inverse of propagation velocity of the waveform above and below the soma along the probe (**Supplementary Fig. 4e-g**). We note that waveform-based quality control metrics did not differ between subdivisions (**Supplementary Fig. 3d**).

Together, these findings show that neurons recorded from MD subdivisions differ across single-channel and multichannel metrics of extracellular waveforms, with MDc neurons showing notably larger amplitudes and more narrow waveforms compared to MDm and MDl neurons.

### Mediodorsal thalamus subdivisions show large differences in spiking metrics

We next examined spiking activity metrics for neurons recorded across MD subdivisions (**Fig. 3a**). Spontaneous firing rate over the entire recording window differed markedly across MD subdivisions (MDm: 5.3 ± 0.2 Hz, MDc: 10.4 ± 0.4 Hz, MDl: 6.4 ± 0.3 Hz, χ^2^= 164.9, *P* < 0.001, Kruskal-Wallis test) with MDc neurons presenting a significantly higher rate compared to MDm and MDl neurons (+98.7%, *P* < 0.001, and +63.7%, *P* < 0.001, respectively, *post-hoc* Dunn’s test for multiple comparisons, **Fig. 3b**). These large firing rate differences across MD subdivisions were similarly evident in passive interval windows (periods outside of bar movement or auditory stimulation related to the behavioral task, **Methods**) (MDm: 4.8 ± 0.2 Hz, MDc: 9.7 ± 0.4 Hz, MDl: 5.5 ± 0.3 Hz^2^ = 143.7, *P* < 0.001, Kruskal-Wallis test), with MDc remaining higher compared to MDm and MDl (+101.1%, *P* < 0.001, and +78.1%, *P* < 0.001, respectively, *post-hoc* Dunn’s test for multiple comparisons, **Fig. 3c**). Indeed, binned firing rates over time were consistently higher in MDc compared to MDm and MDl across the recording period (**Fig. 3d**). Finally, firing rate calculated through an alternative method, using median inter-spike intervals (ISIs), showed a similar pattern of differences across the subdivisions (MDm: 15.5 ± 0.4 Hz, MDc: 22.1 ± 0.6 Hz, MDl: 17.9 ± 0.6 Hz, χ^2^ = 123.4, *P* < 0.001, Kruskal-Wallis test), with MDc higher in rate compared to MDm and MDl (+42.3%, *P* < 0.001, and +23.2%, *P* < 0.001, *post-hoc* Dunn’s test for multiple comparisons, **Fig. 3e**). Thus, across various methods of analysis, MD subdivisions differed in spontaneous firing rates with MDc neurons presenting a consistent and markedly higher rate and MDl neurons showing modestly higher rates compared to MDm neurons.

**Figure 3.**
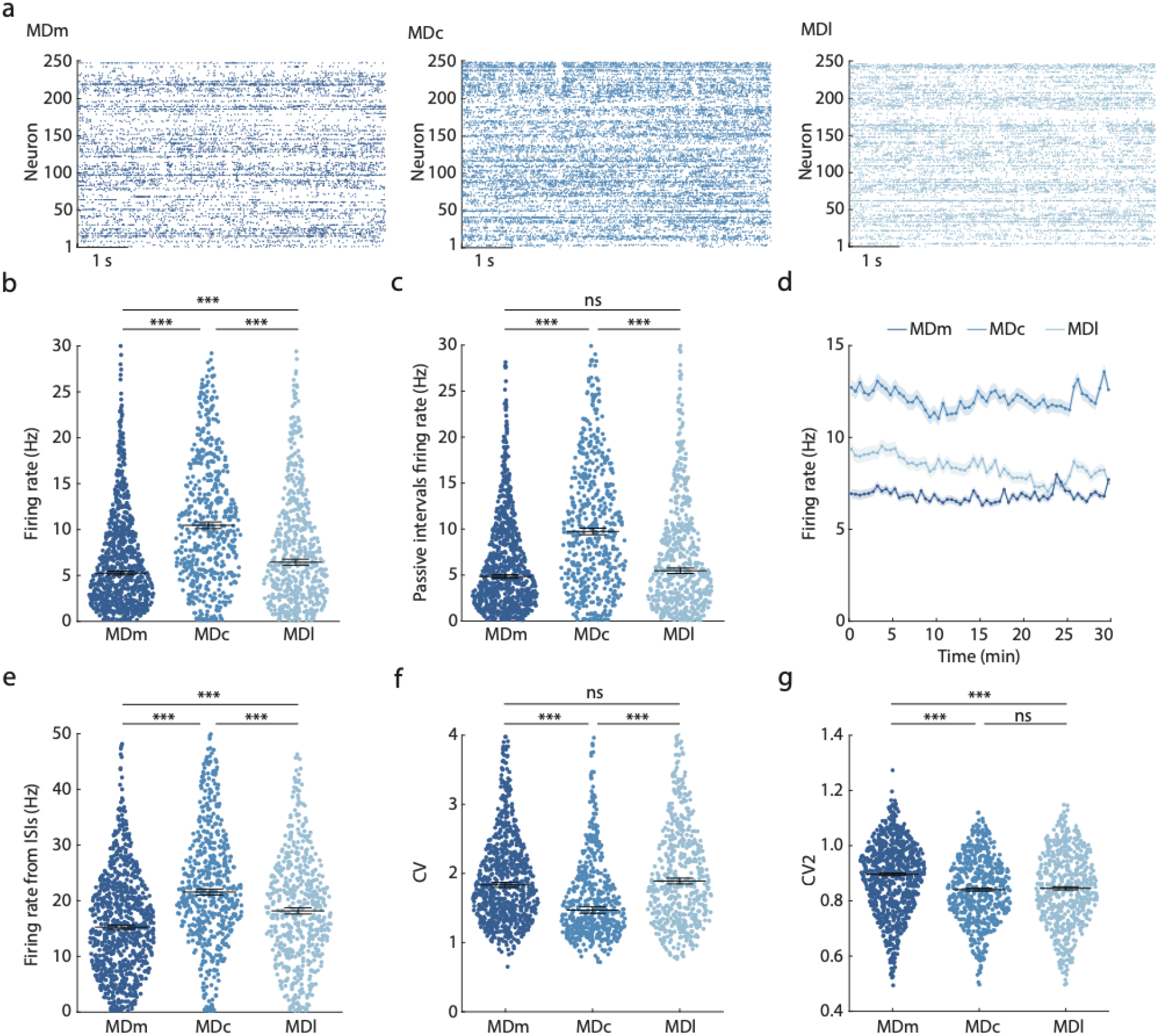
Mediodorsal thalamus subdivisions show differences in spiking metrics. **(a)** Example spike rasters for 250 exemplar neurons in MDm (left), MDc (middle), and MDl (right) over several seconds. **(b)** Swarm plot comparing firing rate for the entire recording period across MD subdivisions, showing differences across subdivisions. Y-axis capped at 30 Hz for clarity. **(c)** Swarm plot comparing firing rate in periods without bar movement or auditory stimulation across MD subdivisions, showing differences across subdivisions. Y-axis capped at 30 Hz for clarity. **(d)** Firing rate across MD subdivisions over time for MDm (dark blue), MDc (blue), and MDl (light blue). Dots indicate mean firing rate per 30-second bins over 60 bins. Shading indicates s.e.m. **(e)** Swarm plot comparing firing rate derived from the inter-spike interval for the full recording period across MD subdivisions, showing differences across subdivisions. Y-axis capped at 50 Hz for clarity. **(f)** Swarm plot comparing CV across MD subdivisions, showing differences across subdivisions. **(g)** Swarm plot comparing CV2 across MD subdivisions, showing differences across subdivisions. Blue dots indicate individual neurons. Black lines in (b), (c), (e), and (f) indicate medians. Black line in (g) indicates means. Error bars in (b), (c), and (e) through (g) and shaded areas in (d) reflect standard error of the means. *** *P* < 0.001; ns non-significant *P* > 0.05. In (b), (c), (e), and (f): Kruskal-Wallis test in followed by *post hoc* Dunn’s for multiple comparisons. In (g): One-way ANOVA test followed by *post hoc* Tukey-Kramer comparisons. Statistics in **Supplementary Table 2.**

We next compared measures of firing rate variability across MD subdivisions. The coefficient of variation of the inter-spike intervals, CV, was different across MD subdivisions (MDm: 1.83 ± 0.04, MDc: 1.47 ± 0.05, MDl: 1.89 ± 0.04, χ^2^ = 104.1, *P* < 0.001, Kruskal-Wallis test), with MDc neurons showing lower CV compared to both MDm and MDl neurons (-19.7%, *P* < 0.001, and -22.3%, *P* < 0.001, *post-hoc* Dunn’s test for multiple comparisons, **Fig. 3f**). Finally, we analyzed CV2, a measure of short-term firing rate variability between consecutive spikes and found modest differences across MD subdivisions (MDm: 0.88 ± 0.01, MDc: 0.84 ± 0.01, MDl: 0.84 ± 0.01, *F* (2, 1927) = 23.9, *P* < 0.001, One-way ANOVA), with MDm neurons showing moderately higher values compared to MDc and MDl neurons (**Fig. 3g**).

Together, these findings suggest that spiking properties of neurons across the three MD subdivisions are different, with most notable increased spontaneous firing rate and decreased rate variability in MDc compared to MDm and MDl.

### Mediodorsal thalamus subdivisions show large differences in burst firing metrics

Burst firing is characterised by transient, high-frequency spiking and is a core feature of thalamic communication^46, 47^. Thus, we next examined differences in burst firing properties of neurons across MD subdivisions. Examination of interspike interval (ISI) histograms revealed a bimodal distribution of ISIs across MD subdivisions (**Fig. 4a**), with global comparison of median ISI duration differing across subdivisions (MDm: 64.4 ± 11.0 ms, MDc: 45.3 ± 19.1 ms, MDl: 55.8 ± 17.4 ms, χ^2^ = 123.4, *P* < 0.001, Kruskal-Wallis test). MDc neurons displayed markedly lower median ISIs compared to both MDm and MDl neurons (-29.7%, *P* < 0.001, and -18.8%, *P* = 0.001, *post-hoc* Dunn’s test for multiple comparisons, **Fig. 4b**). We next fitted a three-component Gaussian mixture model to the ISI distributions of all MD cells, which revealed a threshold at ∼6 ms between the first two Gaussians (**Methods**). We classified bursts as sequences of two or more spikes with consecutive ISIs shorter than 6 ms across the entire recording window.

**Figure 4.**
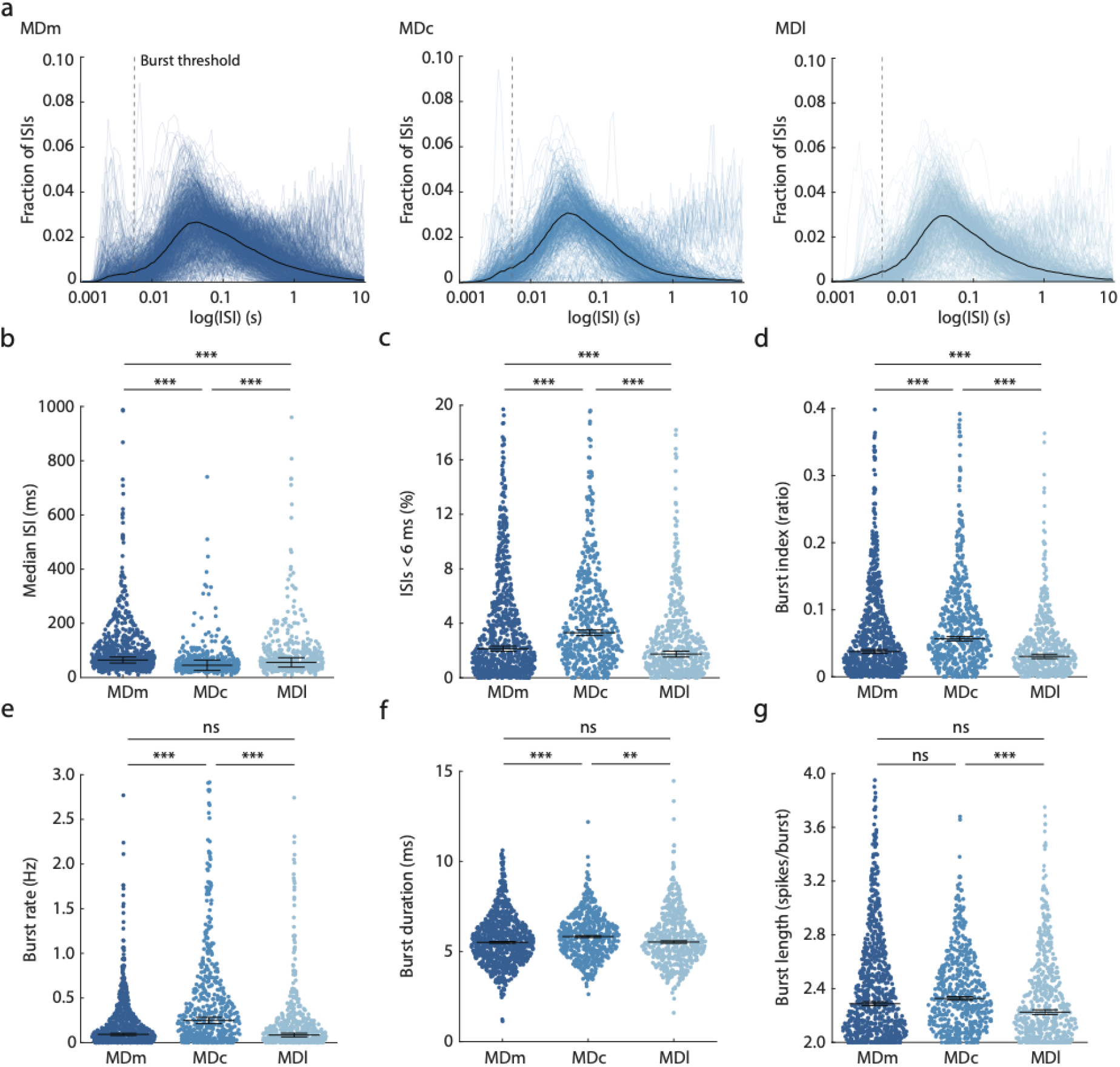
Mediodorsal thalamus subdivisions show differences in burst firing metrics. **(a)** Inter-spike interval (ISI) duration for all individual neurons across MDm (left), MDc (middle), and MDl (right) as a fraction of all neuron spikes. Median ISIs overlaid in black. Dashed vertical grey line indicates the threshold employed for burst firing metrics (**Methods**). **(b)** Swarm plot comparing median ISI duration across MD subdivisions, showing differences across sub-divisions and shortest ISI duration in MDc. **(c)** Swarm plot comparing percentage of ISIs < 6 ms across MD subdivisions, showing differences acros subdivisions. A higher percentage of short ISIs occurs in MDc. **(d)** Swarm plot comparing burst index across MD subdivisions, showing differences across subdivisions and highest burst index in MDc. **(e)** Swarm plot comparing burst rate across MD subdivisions, showing differences across subdivisions and highest burst rate in MDc. **(f)** Swarm plot comparing burst duration in milliseconds across MD subdivisions, showing differences across subdivisions. Longest median burst duration occurs in MDc. **(g)** Swarm plot comparing burst length (number of spikes per burst) across MD subdivisions, showing minor differences across subdivisions. MDl shows shorter burst length compared to MDc and MDm. Blue dots indicate individual neurons. Black lines in (b) through (g) indicate medians. Error bars denote standard error of the means. *** *P* < 0.001; ** *P* < 0.01; ns non-significant *P* > 0.05. Kruskal-Wallis test followed by Dunn’s multiple comparisons. Statistics in **Supplementary Table 2**.

Nearly all neurons in each MD subdivision showed at least one burst in the recording window (MDm: 98.0% [820/837 neurons], MDc: 98.0% [536/547 neurons], MDl: 97.4% [532/546 neurons]). The percentage of ISIs < 6 ms per unit differed across MD subdivisions (MDm: 2.1 ± 0.2 %, MDc: 3.3 ± 0.2 %, MDl: 1.7 ± 0.2 %, χ^2^ = 71.9, *P* < 0.001, Kruskal-Wallis test), with MDc neurons displaying higher percentage of ISIs < 6 ms compared to MDm and MDl neurons (+52.0%, *P* < 0.001, and +90.3%, *P* < 0.001, *post-hoc* Dunn’s test for multiple comparisons, **Fig. 4c**), indicative of increased burst firing rate in MDc. Indeed, burst index, the fraction of spikes with a neighbouring ISI < 6 ms out of all recorded spikes, differed across sub-divisions (MDm: 0.04 ± 0.01, MDc: 0.06 ± 0.01, MDl: 0.03 ± 0.01, χ^2^ = 75.6, *P* < 0.001,

Kruskal-Wallis test), with MDc having a higher burst index compared to MDm and MDl (+51.1%, *P* < 0.001, and +87.1%, *P* < 0.001, *post-hoc* Dunn’s test for multiple comparisons, **Fig. 4d**). Similarly, burst rate differed across subdivisions (MDm: 0.09 ± 0.01 Hz, MDc: 0.25 ± 0.04 Hz, MDl: 0.09 ± 0.02 Hz, χ^2^ = 140.4, *P* < 0.001, η^2^ = 0.07, Kruskal-Wallis test), with MDc neurons presenting a markedly higher burst rate compared to MDm and MDl neurons (+168.6%, *P* < 0.001, and +192.1%, *P* < 0.001, *post-hoc* Dunn’s test for multiple comparisons, **Fig. 4e**). Similar effects on burst index and burst rate differences were obtained when variably classifying bursts as sequences of spikes with consecutive ISIs < 4 ms or < 10 ms (**Supplementary Fig. 5a, b, d, e**). Comparison of burst duration, the average time between the first and last spike within a burst, revealed moderate differences across subdivisions (MDm: 5.51 ± 0.05 ms, MDc: 5.83 ± 0.05 ms, MDl: 5.53 ± 0.07 ms, χ^2^ = 15.7, *P* < 0.001, Kruskal-Wallis test, **Fig. 4f**). Similar differences between the subdivisions were observed for burst length, the average number of spikes per burst (MDm: 2.29 ± 0.02, MDc: 2.33 ± 0.01, MDl: 2.23 ± 0.02, χ^2^ = 16.6, *P* < 0.001, Kruskal-Wallis test, **Fig. 4g**). Notably, this effect was influenced by the ISI threshold used to classify bursts, since MDc showed increased burst length when bursts were classified as sequences of two or more spikes with ISI duration < 10 ms (**Supplementary Fig. 5c, f**). Finally, we found both burst index and burst rate to be moderately correlated with firing rate across all MD subdivisions (*P* < 0.001, Spearman’s rank correlation tests, **Supplementary Fig. 5g-l**), suggesting a positive, not inverse, correlation between spontaneous spiking and burst firing of neurons in MD thalamus.

Together, these findings suggest that MD subdivisions show large differences in burst firing metrics, with prominent higher burst rate of MDc neurons compared to MDm and MDl neurons.

### Robust MD subdivision differences across *in vivo* electrophysiological metrics

Overall, single-neuron *in vivo* electrophysiological recordings in the rodent MD revealed large differences in waveform, spiking, and burst firing metrics across subdivisions, with most prominent differences in MDc neurons in increased extracellular waveform amplitude, higher firing rate, and higher burst rate compared to MDm and MDl neurons. Importantly, the direction and magnitude of these effects were preserved when comparing MD subdivisions with neurons averaged across recording sessions (**Supplementary Fig. 6a-c**; **Supplementary Table 2**), upon re-analyzing the data preceding quality control (**Supplementary Fig. 6d-f**; **Supplementary Table 2**), or upon re-analyzing the data excluding MD cells close to each anatomical border (**Methods**; **Supplementary Fig. 6g-i**; **Supplementary Table 2**). Similarly, the direction and magnitude of the observed effects were also preserved when comparing MD subdivisions for each of the three independently recorded cohorts separately (**Supplementary Fig. 7**; **Supplementary Table 2**).

### The central MD is electrophysiologically distinct from the medial and lateral MD

Considering the cross-metric differences between MD subdivisions, we next examined whether *in vivo* electrophysiological metrics sufficed to classify neuron location within the MD. We trained a linear Support Vector Machine (SVM) based on 15 normalized measures of single-channel and multichannel waveform properties, spiking activity, and burst firing metrics, that differentially correlated to one another across the three subdivisions (**Fig. 5a**; **Methods**; **Supplementary Fig. 8**). The classifier achieved an overall classification accuracy of 52.5% (chance level: ∼33.3%), with MDc yielding the highest performance across subdivisions (∼59.0% accuracy), whereas MDm and MDl were more frequently misclassified as one another (**Fig. 5b**). Training the classifier using exclusively waveform, spiking, or burst firing metrics reduced classification accuracy, but performance remained above chance level (shuffled region labels: ∼33.3%), indicating that each metric category retained some discriminatory information (**Fig. 5c**). Thus, *in vivo* single neuron electrophysiological information incorporating multiple feature classes contained sufficient information to predict anatomical location of neurons within the MD.

**Figure 5.**
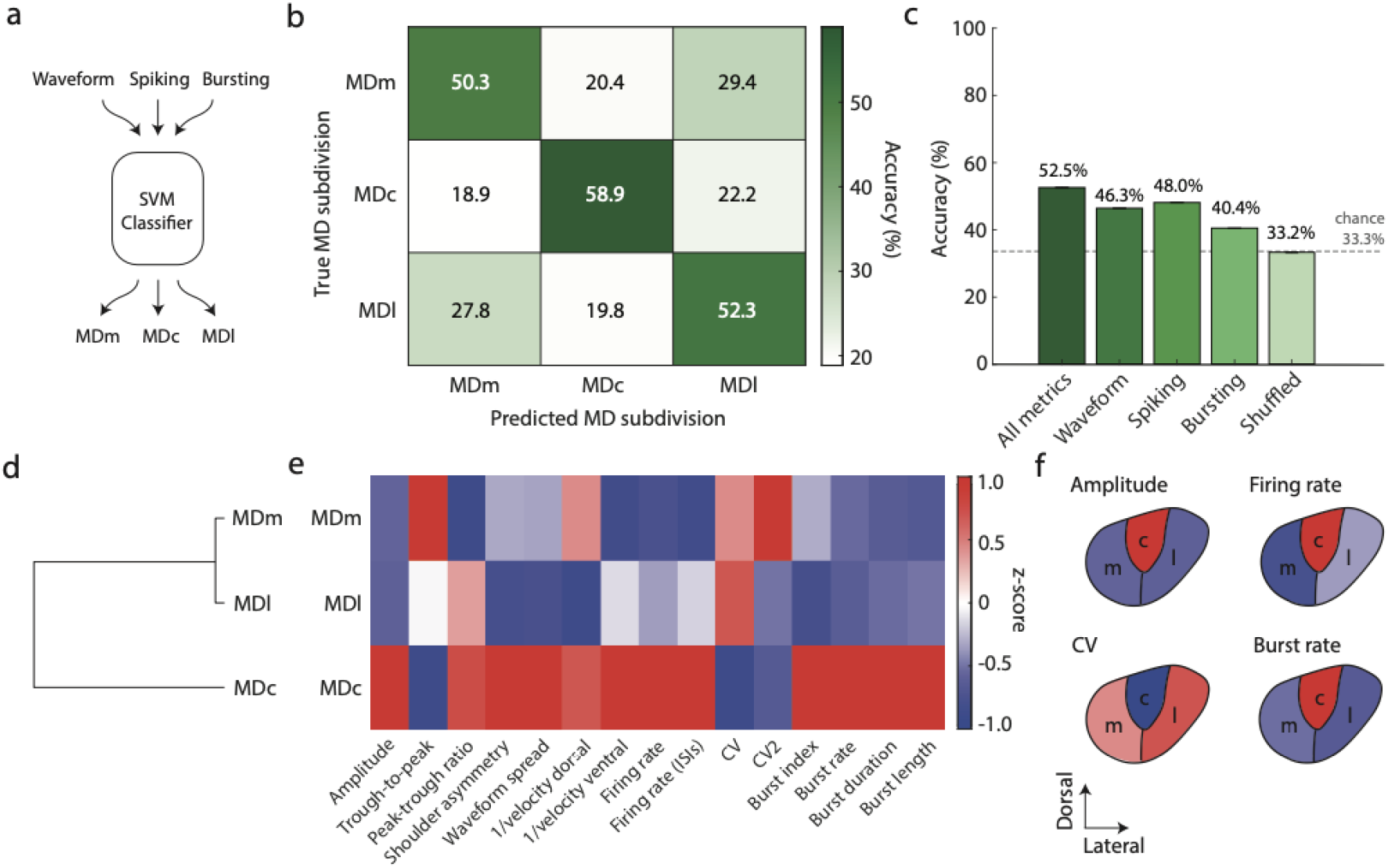
The central MD subdivision is electrophysiologically distinct from the medial and lateral subdivisions. **(a)** Approach. A support vector machine (SVM) classifier was trained to predict MD subdivision location of individual neurons with subsampled equal numbers (*n* = 470 neurons per subdivision) on 15 electrophysiological metrics including waveform, activity, and burst firing. **(b)** Confusion matrix of predicted versus true MD subdivision for neurons across MD subdivisions. Darker green indicates higher accuracy. Chance level is 33%. Classifier accuracy is highest for MDc and lower for MDm and MDl. **(c)** Bar plot comparing prediction accuracy of classifying MD subdivisions based on all electrophysiological metrics (*n* = 15 metrics), only waveform metrics (*n* = 7 metrics, raw single-channel waveform metrics and multichannel waveform metrics), only spiking metrics (*n* = 4 metrics), only bursting metrics (*n* = 4 metrics) and with shuffled MD subdivision labels (*n* = 15 metrics). Bars represent mean classifier accuracy over 100 runs. Dashed line indicates chance level at 33%. **(d)** Hierarchical clustering of MD subdivisions using complete linkage and Spearman’s correlation (ρ) of z-scored median values for 15 electrophysiological metrics. MDm and MDl are more comparable, whereas MDc is more distinct from MDm and MDl. **(e)** Heatmap showing z-scored median values for each electrophysiological metric across MDm, MDc, and MDl. **(f)** Schematic visualization of z-scored median values for the four metrics contributing to the separation of MDc (‘c’) from MDm (‘m’) and MDl (‘l’). CV coefficient of variation, SVM support vector machine.

As MDc was most commonly largely different from MDm and MDl, and classification confusion was higher between MDm and MDl, we next performed hierarchical clustering to investigate the major branches of electrophysiological heterogeneity within the MD. Utilizing the same electrophysiological metrics, hierarchical clustering revealed MDm and MDl to be more similar and clearly distinct from MDc (**Fig. 5d**). Indeed, comparison of normalized electrophysiological metrics revealed MDc differed from MDm and MDl across most metrics (**Fig. 5e**) with normalized amplitude, firing rate, CV, and burst rate contributing most to the separation of MDc neurons from MDm and MDl neurons (**Fig. 5f**). Thus, across multiple features, MDc neurons were electrophysiologically distinct from the comparatively more similar MDm and MDl neurons.

### The central MD clusters with non-MD thalamic nuclei

Differences in waveform features, spiking activity, and burst firing have been sporadically observed as gradients across thalamic nuclei, for example in firstorder versus higher-order nuclei^48^. To compare the distinct electrophysiological profiles of MD subdivisions in relation to other thalamic nuclei, we extended our analysis to include recording sessions from the same animals from thalamic nuclei that are non-MD adjacent, including the posterior (PO), lateroposterior (LP), laterodorsal (LD), and lateral geniculate (LG) nuclei, and the medial geniculate body (MGB) (3 cohorts, 14 animals, 88 recording sessions; **Fig. 6a-d; Methods**). As described above, we employed a combination of validated electrophysiological landmarks to align probe channels with different brain areas (**Supplementary Fig. 9**; **Methods**). Following automated and manual curation of electrophysiological data, we retained a total of 1913 units across these additional thalamic nuclei (PO: *n* = 431, LP: *n* = 221, LD: *n* = 392, LG: *n* = 317, MG: *n* = 552) (**Fig. 6e**; **Supplementary Fig. 10**; **Supplementary Table 3**; **Methods**).

**Figure 6.**
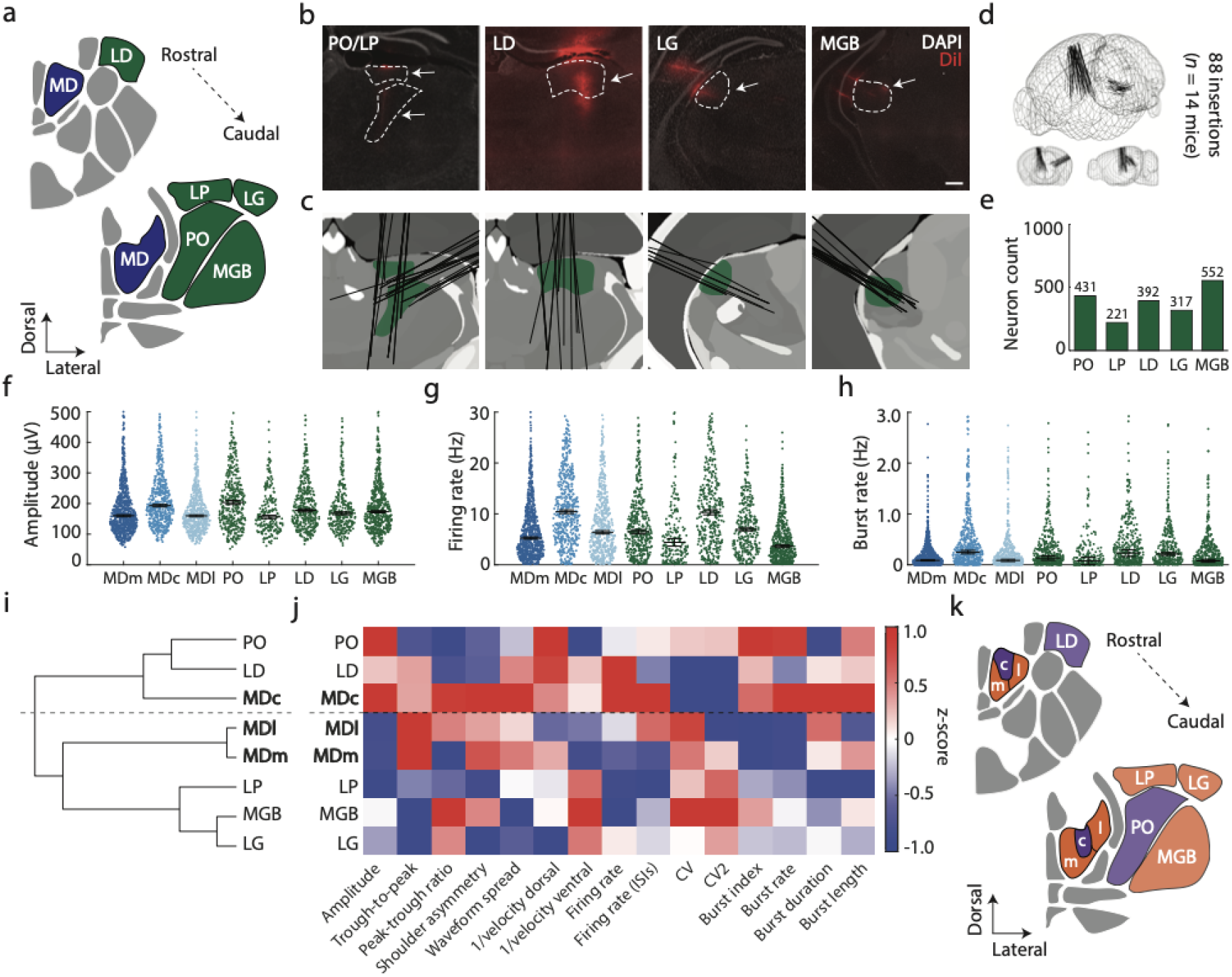
The central MD subdivision clusters with non-MD thalamic nuclei. **(a)** Schematic of additionally recorded thalamic nuclei in coronal view across two anterior-posterior locations. **(b)** Exemplar coronal high magnification epifluorescent microscope image of probe tracks (arrow) through PO/LP, LD, LG, and MGB (outlined by white dashed lines). DAPI nuclear marker in white, DiI fluorescence probe marker in red. Scale bar, 200 μm. **(c)** All probe insertions (black lines) through PO/LP, LD, LG, and MG (regions highlighted in green). Details on probe channel realignment to brain regions in **Supplementary Fig. 9**. **(d)** 3D schematic of brains showing all reconstructed probe trajectories (black lines) for PO, LP, LD, LG, and MGB. **(e)** Total number of recorded neurons for each thalamic nucleus following quality control and manual curation. Details on quality control in **Supplementary Fig. 10**. **(f)** Swarm plot comparing waveform amplitude across MD (blue) and non-MD (green) thalamic nuclei. Y-axis capped at 500 µV for clarity. **(g)** Swarm plot comparing firing rate across MD (blue) and non-MD (green) thalamic nuclei. Y-axis capped at 30 Hz for clarity. **(h)** Swarm plot comparing burst rate across MD (blue) and non-MD (green) thalamic nuclei. **(i)** Hierarchical clustering of MD and non-MD thalamic nuclei subdivisions using complete linkage and Spearman’s correlation (ρ) of z-scored median values for 15 electrophysiological metrics. MDm and MDl cluster together, while MDc clusters with other non-MD nuclei. **(j)** Heatmap showing z-scored median values for each electrophysiological metric across thalamic nuclei. **(k)** Schematic visualization of clustering of thalamic nuclei. MDc clusters more closely with LD and PO on physiology (purple), while MDl, MDm, LP, MGB, and LG cluster more closely together (orange). Dots indicate individual neurons. Black lines in (f) through (h) indicate medians. Error bars denote standard error of the means. PO posterior nucleus of the thalamus, LP lateral posterior nucleus of the thalamus, LD laterodorsal nucleus of the thalamus, MGB medial geniculate body of the thalamus.

As waveform amplitude, firing rate, and burst rate were the electrophysiological metrics most prominently different across MD subdivisions, we first examined these across all thalamic nuclei included. We observed substantial heterogeneity across thalamic nuclei for all metrics, with nuclei showing distinct profiles particularly for firing rate and burst rate (**Fig. 6f-h, Supplementary Table 2**). Hierarchical clustering for all recorded nuclei including MD based on 15 single- and multichannel waveform features, spiking activity metrics, and burst firing characteristics (**Methods**) revealed a clear separation between MDc and the other MD subdivisions: MDm and MDl formed the earliest cluster and subsequently grouped with LG, MGB, and LP. In contrast, MDc formed a cluster with PO and LD before merging with the remaining thalamic nuclei including MDm and MDl (**Fig. 6i**). Comparison of normalized electrophysiological metrics revealed widespread and non-uniform differences across all thalamic nuclei (**Fig. 6j**). Notably, the MDc, PO, LD cluster showed relatively larger waveform amplitudes, higher firing rates, and increased bursting metrics as compared to the MDl, MDm, LP, LG, and MGB cluster. Thus, the two major clusters identified (**Fig. 6k**) are differentiated across multiple waveform, spiking, and bursting metrics.

Together, these findings indicate that MDc shares greater electrophysiological similarity with other non-MD thalamic nuclei and is distinct from the rest of the MD.

## DISCUSSION

The MD nucleus of the thalamus is implicated in a broad range of cognitive processes and in the pathophysiology of neurological and psychiatric brain disorders. Here, we performed high-density single-neuron *in vivo* electrophysiological recordings with Neuropixels probes across the MD of head-fixed, behaving wildtype male mice. We *post-hoc* assigned simultaneously recorded single neurons to MDm, MDc, and MDl to allow examination of electrophysiological differences across MD subdivisions. Our analyses revealed MD subdivisions to be strikingly different across single- and multichannel waveform, spiking activity, and burst firing metrics. These effects were consistently observed across independent cohorts, recording configurations, present throughout recording duration, and independent of quality control or exact subdivision anatomical border location. Thus, our findings show that MD subdivisions display distinct *in vivo* electrophysiological profiles.

Our findings increase the dimensionality of differences between MD subdivisions. MDm, MDc, and MDl are classically shown to have distinct anatomical input-output connectivity patterns^1, 29^ and were more recently shown to have different gene expression profiles^34, 41, 42^ and *ex vivo* electrophysiology^31, 33, 35^. In line with these previous findings, we show that MD subdivision differences are also prominent at the level of *in vivo* electrophysiology in awake, behaving animals. Neurons across MD subdivisions were shown to have non-uniform electrophysiological profiles, with MDc dissimilarity evident from various electrophysiological measures, suggesting that the combination of waveform, spiking, and bursting features distinguishes MD subdivisions. Previously, transcriptomic characterization suggested a gradient of gene expression across and within thalamic nuclei, from midline nuclei to lateral, sensory nuclei^42^. This mediolateral gene expression gradient was accompanied by continuous variation in cortical projection profiles, morphology, and *ex vivo* electrophysiological properties^42^. Indeed, multiple studies across animal models and humans have provided evidence for the presence of gene expression, cytoarchitecture, and thalamocortical connectivity gradients across and within thalamic nuclei^42, 49–52^. The current findings indicate that MD subdivisions do not adhere to a continuous mediolateral gradient of *in vivo* electrophysiological properties. Together, these findings support the emerging notion that thalamic organization is governed by multiple discrete and continuous features that potentially vary along non-overlapping anatomical axes^49^. Overall, our findings contribute and further support the increasingly acknowledged notion that MD subdivisions could represent distinct populations of neurons with divergent connectivity, physiology, and functionality.

The various sources of heterogeneity across MD subdivisions are likely mechanistically linked. Divergent expression profiles of ion channels and other synaptic proteins within each MD subdivision may underlie distinct basal waveform properties as well as divergent spiking and bursting characteristics. Previously, transcriptomic and *ex vivo* electrophysiological characterization of thalamic nuclei suggested MDc belongs to a different molecular profile than MDm and MDl^42^. Neurons within the MDc molecular profile were shown to have smaller action potential width^42^, which is in line with the shorter trough-to-peak latencies for MDc neurons reported here. We highlight the presence of larger, more narrow waveforms in MDc (**Fig. 2**), which throughout the brain often characterize faster-spiking neurons^53, 54^. Higher *in vivo* burst firing in MDc (**Fig. 4**) aligns with findings from a recent preprint which reported increased sag in membrane voltage *ex vivo* and increased hyperpolarization-activated cyclic nucleotide-gated (HCN) channel activity in MDc neurons^31^. However, increased HCN activity correlated with lower input resistance and decreased action potential firing, suggestive of MDc neurons being less excitable than MDm and MDl neurons^31^. Here, we find MDc to have remarkably increased firing rate compared to MDm and MDl (**Fig. 3**). *In vivo* electrophysiological differences in firing rate could in part reflect distinct patterns of afferent connectivity of MD subdivisions with cortical and subcortical partners which are minimally preserved in *ex vivo* electrophysiology^1, 29^. Conversely, differences in firing rate and bursting activity could be attributed to differential inhibitory control of MD subdivisions from thalamic reticular nucleus (TRN)^55^ or local GABAergic interneurons. Whether increased activity in MDc reflects increased intrinsic excitability, increased synaptic inputs, or reduced inhibition, remains to be determined. Overall, differences at the levels of anatomical input-output connectivity, molecular expression, *ex vivo* and *in vivo* electrophysiology converge to suggest that heterogeneity across the MD may share common underlying mechanisms.

Hierarchical clustering within the MD and across other thalamic nuclei revealed that MDc neurons surprisingly shared greater electrophysiological similarity with neurons in PO and LD compared to neurons in MDm and MDl. Briefly, PO is a higher-order thalamic nucleus involved in sensorimotor processing which shows widespread connectivity with sensory and motor cortical areas^56, 57^. LD is a higher-order thalamic nucleus involved in spatial learning and memory which is interconnected with the anterior cingulate and retrosplenial cortices^58^. We note that while MDc clustered exclusively with higher-order thalamic nuclei, MDm and MDl primarily clustered with first-order thalamic nuclei (LG and MGB)^42^. MDc, PO, and LD neurons had relatively larger waveform amplitudes, higher firing rates, and higher burst indices than neurons across the remaining thalamic nuclei (**Fig. 6**). These findings are partially in line with previous transcriptomic findings that show MD, PO, and LD cluster closely along the axis of gene expression variation in the thalamus^42^. Thus, the observed electrophysiological similarities between MDc, PO, and LD neurons could in part reflect overlapping gene expression profiles of ion channels and neurotransmitter receptors. In parallel, electrophysiological similarities could in part be driven by shared cortical and subcortical inputs to MDc, PO, and LD. Overall, these findings further reinforce the notion that MDc neurons represent a physiologically distinct population within the MD.

The current findings represent the largest single-neuron *in vivo* electrophysiological interrogation of the rodent MD to date. However, several considerations remain. First, *in vivo* electrophysiological recordings were obtained unilaterally in right hemisphere and therefore potential hemispheric differences across MD subdivisions remain to be assessed. Second, electrophysiological recordings were performed exclusively under head-fixation and during a sensorimotor behavioral task and may not entirely capture the functional dynamics of MD neurons. How the observed basal electrophysiological differences facilitate distinct computational demands of sub-division-specific thalamocortical communication remains an important open question. For example, narrow waveforms, higher firing rates, and increased burst firing may influence how MDc neurons encode and relay information to their cortical and subcortical partners. Thus, further investigations should focus on identifying how differences in electrophysiological metrics across MD subdivisions influence behavioral paradigms^3, 5, 7, 8^. Additionally, we did not record during any sleep stages, which could illuminate further subdivision differences in electrophysiology. Similarly, whether additional characteristics such as local field potential measures or spike locking to local oscillations differentiate between subdivisions remains to be determined. Moreover, recordings were conducted in male mice, limiting generalizability of the findings to females where sexual dimorphism in thalamic physiology, in addition to previously reported thalamic size^59^, may exist. Finally, considering established interspecies differences in MD anatomical connectivity, cytoarchitecture, and molecular expression^34, 48, 60^, the relevance of these *in vivo* electrophysiological profiles of MD subdivisions in other species, particularly in humans, remains to be investigated. Thus, the full extent of electrophysiological differences across MD subdivisions during naturalistic behaviors, cognitive tasks, and during sleep remains to be elucidated.

Nonetheless, our findings pose important implications for future studies involving MD in both health and disease. First, given profound physiology differences, rodent studies examining MD function should employ investigations at the level of subdivisions to allow characterization of potential functional and pathological diversity within the MD. Second, while cross-species homology remains uncertain, human neuroimaging studies could incorporate MD subdivision-segmentation to determine whether subdivisions differentially engage in cognitive functions and/or are selectively implicated in the pathophysiology of different disorders. Third, future state-of-the-art neuromodulation approaches targeting the MD in disease including deep brain stimulation and transcranial focused ultrasound may require subdivision-specific spatial resolution, as divergent electrophysiological properties of MD subdivisions could shape responsiveness to stimulation and thus influence downstream effects and therapeutic efficacy. Multi-feature classification as described here could be employed for optimized online targeting of stimulation leads in MD thalamus. Overall, improved physiological and functional characterization of MD subdivisions could improve understanding of pathology involving thalamus, support brain stimulation interventions in human patients, and ultimately guide therapeutic strategies.

Here, we present *in vivo* electrophysiological findings that extend the parameter space of differences across MD subdivisions. We report that MDm, MDc, and MDl show divergent *in vivo* electrophysiological profiles, with MDc most distinct from MDm and MDl across various metrics. These findings pose implications both for examination of MD functioning in healthy behavior in humans and animal models as well as for potential therapeutic targeting of the MD in neurological and psychiatric disorders.

## RESOURCE AVAILABILITY

## Lead contact

Requests for further information and resources should be directed to and will be fulfilled by the lead contact, Jeffrey Stedehouder.

## Material availability

This study did not generate new unique reagents.

## Data and code availability

All data supporting the figures are available in the manuscript or supplementary materials. Processed data and core Matlab analysis code will be made available through the Medical Research Council (MRC) Centre of Research Excellence in Restorative Neural Dynamics and Brain Network Dynamics Unit (BNDU) data sharing platform upon publication (https://data.mrc.ox.ac.uk/mrcbndu/data-sets/search). Any additional information required will be made available from the lead contact upon request.

## ACKNOWLEDGMENTS

This research was supported by a Medical Research Council Oxford Doctoral Training Partnership Scholarship (MR/W006731/1) and an Onassis Foundation Scholarship (F ZU 056-1/ 2024-2025) to KP, a Medical Research Council UK award (MC_UU_00003/6) to AS, a Sir Henry Wellcome Postdoctoral Fellowship (224129/Z/21/Z), John Fell Fund Oxford award (0013599) and Win Seed Grant to JS, and a Medical Research Council award (UKRI/MR/B000936/1). The authors would like to thank Jane Westcott, Rae Dolman, and Alisha Miller for animal technical assistance and Ben Micklem for IT support.

## AUTHOR CONTRIBUTIONS

**Katerina Panti**: Conceptualization, Methodology, Investigation, Visualization, Writing – original draft, Writing – review & editing. **Charlotte J. Stagg**: Writing – review & editing, Funding acquisition. **Andrew Sharott**: Writing – review & editing, Funding acquisition, Supervision. **Jeffrey Stedehouder**: Conceptualization, Methodology, Investigation, Visualization, Writing – review & editing, Funding acquisition, Supervision.

## SUPPLEMENTAL INFORMATION

Document S1. Figures S1-S10 and Tables S1 and S3

Table S2. Statistics

## MATERIALS AND METHODS

### Mice

All experiments were conducted according to the UK Animal Scientific Procedures Act (1986) and under personal and project licenses released by the Home Office following appropriate ethics review. In total, 14 C57BL/6J adult male mice ranging in age from 12 to 20 weeks were used. Four C57BL/6J wildtype male were directly obtained from Charles River Laboratories (‘Cohort 1’). Six wildtype mice were obtained from breedings between male heterozygous *Grin2a*^+/-^, backcrossed into C57BL/6J background and female wildtype C57BL/6J mice obtained from Charles River (‘Cohort 2’). Data for Cohort 2 has been previously acquired and described elsewhere^12^. Four wildtype mice were F1 offspring obtained from breedings between male *Sp4*^+/-^in 129Sv background and female wildtype C57BL/6J mice obtained from Charles River (‘Cohort 3’; **Supplementary Fig. 1**). Mice were group-housed in individually ventilated cages on a 12-hour light/dark cycle pre-surgery with constant temperature (∼22°C) and single-housed post headplate surgery. Food pellets were provided *ad libitum*. Mice had controlled access to water during habituation, training, and recording.

### Surgery

Headplate surgery for all cohorts was performed as described previously^12^. Briefly, mice were anaesthetized with isoflurane (induction at ∼4% and maintenance at ∼1.5-2%) in O_2_ (∼1 L/min). Following induction of anesthesia, analgesia was provided using Vetergesic (0.08 mg/kg in sterile saline) administered subcutaneously as well as with local Marcaine application (8 mg/kg). The skin and periosteum on the dorsal surface of the skull was removed, and the surrounding skin secured to the skull using cyanoacrylate (Vetbond, 3M, Minnesota, USA). The skull was prepared using 3% H_2_O_2_ and etched using a bone scraper. A custom-made titanium headplate was implanted on each animal (Get It Made Ltd, London; 0.7 g, internal diameter 9 mm), aligned to lambda and the midline and secured using dental cement (Super Bond C&B, Parkell, United Kingdom). An M1 3-mm reference screw (Precision Technology, East Grinstead) was wrapped in silver wire and placed into a craniotomy over right cerebellum and fixed in place using dental cement (Jet Denture Repair Powder: Lang Dental, Illinois, USA; Meadway Repair Liquid: MR. Dental, Surrey, United Kingdom). A custom-designed, 3D-printed shield was secured on top of the headplate. Mice were allowed to recover for 7 days post-surgery, followed by habituation to head-fixation and behavioral training.

Twenty-eight days after headplate surgery, anesthetized recovery surgery was performed to create small craniotomies (∼0.5-1 mm) above each recording site. For thalamus recordings, anterior and posterior craniotomies were performed with coordinates and planned craniotomy sites that differed slightly for each of the three cohorts. Following surgery, the craniotomies were covered with DuraGel (Cambridge Neurotech, United Kingdom) and overlaid with silicon (Body Double, Smooth-On, United Kingdom).

Anterior craniotomy over the thalamus (mm from Bregma):

Cohort 1: AP -2.00, ML +1.44

Cohort 2: AP -1.00, ML +1.20

Cohort 3: AP -1.00, ML +1.00.

Posterior craniotomy over the thalamus (mm from Bregma):

Cohort 1: AP -2.50, ML -4.5

Cohort 2: AP -2.5, ML -4.70

Cohort 3: AP -2.50, ML -4.70.

### Behavioral task

The lever pressing task has been described in detail elsewhere^12^ with minor variations between cohorts. Briefly, mice were habituated to head-fixation for ∼7 days in incremental durations before the onset of behavioral training. During training, mice had controlled access to water and were maintained at ∼85 - 87% of their *ad libitum* body weight. Mice were placed in a tube with custom-made interior lining with forepaws resting on two metal bars under head-fixation. A moveable waterspout was positioned ventral and anterior to the mouse within licking distance. A PyControl system ^61^ was used to control dispensation of reward (5% (wt / vl) sucrose in water, ∼5 μl) droplets through the spot.

Mice were trained to push the left metal bar self-paced forward over a distance of ∼5-10 mm and return it to starting position. All bar pushes were coupled to a pure tone (2-16 kHz, 100 ms, ∼75 dB) delivered from a speaker (OpenEphys) to the right side of the mouse at a distance of ∼15 cm. Only bar pushes that were returned to starting position within 250-1000 ms were labelled as correct and rewarded. On correct trials, reward was delivered 1.5 seconds after the bar was returned to home position. Mice were trained on ∼20-35 min sessions once per day for two weeks (mean: 11.75 sessions; range 11 – 13 sessions).

During recordings, mice were exposed to the same bar pushing paradigm, with small changes and divided into three stages: In stage (1), mice performed 30 standard trials similar as during training. In stage (2), mice performed up to 500 trials or trials for 30 mins, whichever was attained earlier. During this stage, bar presses could elicit a tone of 1 octave different from the training tone on a subset (15%) of trials. In another 15% of trials, bar presses did not elicit any tone. In addition, every ∼20-40 seconds, the target tone was played passively in absent of any bar press. All remaining trials were as in stage (1). In stage (3), the bar was locked in place, and mice were passively exposed to a series of 30 tones across 4 frequencies with an interval of ∼ 2 seconds (totalling 120 tones). Further details on the task and behavioral performance can be found elsewhere^12^.

### Probe trajectory planning

Probe trajectories were planned using the Allen Institute Common Coordinate Framework (CCF) GUI tool^62^ and PinPoint^63^ with the following estimated coordinates:

Anterior thalamus probe (mm from Bregma):

Cohort 1: -2.00-2.75 AP, +1.25-1.75 ML, 10-15° ML tilt, 0° AP tilt

Cohort 2: -1.00-1.75 AP, +1.00-1.50 ML, 0-7° ML tilt, 0-10° AP tilt

Cohort 3: -1.00-1.75 AP, +0.80-1.30 ML, 5-10° ML tilt, 0° AP tilt

Posterior thalamus probe (mm from Bregma):

Cohort 1: -2.30-2.70 AP, -4.70-4.20 ML, 70° ML tilt, -10-12° AP tilt

Cohort 2: -2.30-2.70 AP, -4.90-4.40 ML, 60-70° ML tilt, -10-15° AP tilt

Cohort 3: -2.30-2.70 AP, -4.90-4.40 ML, 55-65° ML tilt, -10-15° AP tilt

### Electrophysiological recordings

Recording details are provided elsewhere^12^. Up to four single-shank Neuropixels 1.0 probes (IMEC, Belgium) were used to record activity simultaneously from cortex, hippocampus, thalamus, and striatum acutely during rest and during performance on the bar pressing task. Electrophysiological recordings were made using OpenEphys software^64^, with default settings used for gain (AP band 500x, LFP band 250x), sampling rate (spike band 30 kHz, LFP band 2.5 kHz), and channel mapping (384 electrodes from tip). Here, we include and analyse recording from mediodorsal thalamus (MD), and additionally laterodorsal thalamus (LD), lateral geniculate nucleus (LG), medial geniculate body (MGB), lateroposterior nucleus (LP), and posterior nucleus (PO). Neurons from MD, LD, LP, and PO were recorded from the right hemisphere (anterior thalamus probe). Neurons from LG, LD, and MGB were recorded from the left hemisphere (posterior thalamus probe). Custom 3D printed probe holders and headstage holders (10k resin, Formlabs, United Kingdom) were used to support probes and headstages, respectively. Probes were lowered using individual automated single-axis motors (700-105140-IVM, Scientifica, United Kingdom) mounted on a custom-designed frame with Kopf micromanipulators (DKI 1760-61-SB) were lowered with a speed of 4-5 μm/s for the first ∼2000 μm, followed by a speed of 3 μm/s till recording depth. Upon reaching target location, probes were retracted by 100 μm at a speed of 3 μm/s and allowed to settle in place for ∼5 mins before recording. Probe recordings were referenced to the probe tip. A small amount of sterile saline was applied to the craniotomy during probe lowering, recording and retraction. After recording, probes were slowly retracted at a speed of 5-6 μm/s. At the end of a recording session, the craniotomy was again covered with DuraGel and overlaid with silicone. After recording, probes were cleaned in 1% tergazyme, de-ionised water, and isopropanol.

Electrophysiological recordings were made using OpenEphys software^64^ at 30 kHz with default parameters. Each recording session lasted ∼34.5 minutes (range 29.7 – 38.9 mins), with an average total head-fixation duration of < 120 mins. Recording sessions occurred once per day per mouse for ∼5 days (range 4 – 7 days). PyControl was used to acquire event times^61^.

Neuropixels probes were sharpened at ∼45° angles using a repurposed hard drive disk. The ground and reference pads of the probes were shortened and soldered to a wire that connected to the animal and ground. Each probe was labelled with a fluorescent lipophilic membrane stain 1,1’-dioctadecyl-3,3,3’,3’ - tetramethylcarbocyanine perchlorate (DiI; Invitrogen, California, USA; ∼1-2 mg/ml in isopropanol) ∼5 minutes before each animal was head-fixed and ∼15 minutes before probe insertion. Time from DiI labelling to probe insertion did not differ across cohorts.

### Histology

One day after the last recording, mice received terminal anesthesia with a pentobarbital intraperitoneal injection (200 mg/kg). Mice were transcardially perfused with 0.1 M phosphate buffer (PB, pH 7.4; ∼10 ml) followed by 4% paraformaldehyde solution (PFA) in PB (∼50-75 ml). Brains were removed and post-fixed for 2 hours in 4% PFA at room temperature, washed with 0.1 M PB and placed overnight in 10% sucrose/0.1 M PB at 4°C. Brains were embedded in 12% gelatin/10% sucrose using custom, 3d-printed embedding wells, post-fixed in 10% PFA/30% sucrose for 2 hours and stored in 30% sucrose at 4°C until cutting. Whole brains were coronally sectioned into 50 µm slices using a sliding microtome (Expredia HM450, Fisher Scientific, United Kingdom), with slices sequentially collected in 0.1 M PB. Slices were immersed in 1:5000 DAPI nuclear marker for ∼5 mins, mounted on glass slides, and coverslipped using Vectashield mounting medium (H-1000).

Epifluorescent images for both DAPI and DiI were acquired using an Axio Imager M2 epifluorescent microscope (Zeiss), equipped with Plan-Achromat objective lenses, a Hamatsu Flash 4.0 LT camera (C10600) and Colibri LEDs. ZEN Blue software was used to acquire tiled single plane images of each entire slice with a 10x objective.

Probe tracks were reconstructed using the Allen Institute Common Coordinate Framework (CCF) GUI Matlab tool, SHARP-Track^65^. Coronal images including the entire brain except the cerebellum were processed and aligned to the matching coordinates on the reference atlas, with probe tracks manually traced on the DiI fluorescence (∼20 probe points per track), with the deepest probe track point as the most ventrally recovered fluorescent signal. Probe track data were utilized to map brain areas to channels along the length of the Neuropixels probe and imported into CellExplorer environment^66^.

### Regional channel alignment

To align channels from reconstructed tracks to brain regions, we utilized a combination of neuron count as well as single neuron and local field potential (LFP) metrics for realignment of probe channels with electrophysiology using a custom-written GUI. For the MD, probes dorsoventrally traversed cortex, hippocampus, thalamus, and hypothalamus and channels were aligned to brain regions using previously validated landmarks, sequentially including brain surface (no units, uniform LFP power), white matter border from cortex to hippocampus (no units, uniform LFP power), CA1 pyramidal cell layer (high unit density and LFP phase reversal), dentate gyrus (high unit density, peak LFP power), transition border from hippocampus to thalamus (no units, lower LFP power), and transition border from thalamus to hypothalamus (lower units, activity)^67–70^. The channels between these borders or landmarks were linearly rearranged to fit brain region identity to reach channel. Thus, MD targeting was a result of histological information (deepest fluorescent signal) and ∼4-5 electrophysiological landmarks across the full length of the probe. Similar electrophysiology landmarks were used to align channels to brain areas for probe tracks recording from other thalamic nuclei (LP, PO, LD, LG, and MGB). Although not included in the realignment, we note that lateral habenula showed high spiking, evident in the action potential band activity.

### MD subdivision assignment

To assign channels of the MD to the three MD subdivisions, we updated the Allen Institute Common Coordinate Framework (CCF) Atlas^62^ with coordinates from Paxinos and Franklin^71^ for MD subdivisions (**Supplementary Fig. 2**). In the Paxinos and Franklin atlas, thalamic nuclei have been described based on previous work.^71, 72^ In general, the central subdivision (MDc) was shown to be richest in myelinated fibers and was thus easily segregated from the medial (MDm) and lateral (MDl) subdivisions. Additionally, MDl had the highest activity for acetylcholinesterase^72^. The Paxinos and Franklin atlas has been used to delineate MD subdivisions in previous studies^31^. Here, we divided the MD across its mediolateral axis into MDm, MDc, and MDl, symmetrically in both hemispheres. The most anterior and posterior sections of the MD (anterior to -1.00 mm bregma and posterior to -2.00 mm bregma) were not included, as these contain no clear segregation into subdivisions in Paxinos and Franklin^71^. Neurons assigned to the most anterior or posterior regions of the MD were removed from further analysis (< 10%). In an additional analysis (**Supplementary Fig. 6g-i**), we removed all neurons located with 100 µm of an anatomical border (either between subdivisions or with other brain regions), to exclude effects of region border alignment variation.

### Quality control

Automated spike sorting of electrophysiological data probes was completed with Kilosort 3.0 using default parameters^73^. Neurons located to MDm, MDc, and MDl labelled as ‘good’ by Kilosort3 were selected for downstream analyses. Non-somatic spikes were removed with custom-written Matlab scripts, removing single-channel waveforms that either contained a first peak with an amplitude larger than the main waveform trough, or when the second peak following the trough was larger in amplitude than the main waveform trough. Further quality control was performed utilizing a combination of previously described ^67, 70, 74^ spiking and waveform metrics including neurons with presence ratio (>0.7), total spike count (>300), and average waveform amplitude (>20 μV), filtered using BombCell^75^ and custom-written Matlab scripts.

For the MD, resulting neurons (*n* = 2110) from 14 animals were further individually manually curated using a custom-written Matlab GUI by two independent raters (K.P. and J.S.). In total, another 180 MD neurons were removed following manual curation, a similar number between MD subunits (MDm: 87 neurons (∼9.4%), MDc: 46 neurons (∼7.8%), MDl: 47 neurons (∼7.9%)). Overall, comparable numbers of neurons were discarded across the 3 subdivisions: (MDm: 200 neurons (∼19.3%), MDc: 204 neurons (∼27.2%), MDl: 99 neurons (∼15.3%)) (**Supplementary Fig. 3**). The final curated dataset consisted of 1930 MD neurons (MDm: 847 neurons, MDc: 547 neurons, MDl: 546 neurons) across 12 animals, which were used for all subsequent analyses

For other thalamic nuclei (including LD, LG, MGB, LP, PO), resulting neurons following automated curation (*n* = 2382) from 14 animals were similarly manually curated using a custom-written Matlab GUI by two independent raters (K.P. and J.S.). In total, 469 neurons were removed across these thalamic nuclei following manual curation, a similar number across all nuclei (LD: 26 neurons (∼5.6%), LG: 46 neurons (∼11.2%), MG: 24 neurons (∼5.0%), LP: 21 neurons (∼6.8%), PO: 25 neurons (∼4.9%)) ((**Supplementary Fig. 10**).The final curated dataset consisted of 1913 units across these additional thalamic nuclei (PO: *n* = 431, LP: *n* = 221, LD: *n* = 392, LG: *n* = 317, MG: *n* = 552).

### Analysis of waveform metrics

Neural waveform metrics were calculated across the full recording duration unless otherwise specified. Raw waveforms consisted of 48 data points and using a quadratic interpolation in Matlab v2021b were increased 100-fold to 4800 data points^76^. Single-channel extracellular waveform features were calculated from these interpolated raw waveforms (non-filtered, ‘refined’ waveforms) across the full recording duration. The probe channel where the mean waveform amplitude was largest (‘best channel’) was used for analysis of single-channel metrics and these included: amplitude (in μV) was calculated as the difference between the maximum negative deflection and the maximum positive deflection of the waveform. Trough-to-peak latencies, or repolarization latencies, (in ms) were calculated by subtracting the trough negative deflection latency from the peak positive deflection latency following the trough. Peak-trough ratio was calculated as the ratio between the amplitude of the first peak following the trough and the amplitude of the trough negative deflection. Shoulder asymmetry was calculated as the ratio between the positive peak preceding the trough and the positive peak following the trough. Single-channel waveform metrics were additionally re-examined using filtered (500 Hz high-pass filter) waveforms interpolated in a similar manner (**Supplementary Fig. 4a-e**).

Furthermore, we analysed multichannel extracellular waveform features by considering waveforms of a single sorted unit recorded from multiple adjacent recording channels as a multichannel waveform^45^. Only the side of the best channel was used to calculate multichannel waveforms, with the vertical distance between probe channels set at 20 μm. For multichannel waveform analysis, we extracted three features in the spatial axis: Spread along the probe (‘waveform spread’), and the inverse of propagation velocity above and below the soma along the probe (‘1/velocity dorsal’ and ‘1/velocity ventral’, respectively).

The distribution of the waveform amplitude across channels was used to define the waveform spread of each unit. Amplitude was plotted against probe channel distance from the best channel, resulting in a curve with peak at t = 0. Waveform spread was defined as the range of channels along the probe (max 12 channels away from the best channel on the vertical axis) with amplitude >12% of the maximum amplitude. For information on signal propagation velocity, we calculated the inverse of velocity (ms/mm) by fitting two regression lines to the time of waveform trough for each channel against the distance of the channels relative to the best channel^77^. As reported previously for other thalamic nuclei and different from cortical principal neurons^45^, dorsal and ventral propagations of the waveforms in MD thalamus neurons were highly symmetrical across each subdivision, in line with their multipolar dendrite morphology^37^.

### Analysis of spiking activity metrics

Spontaneous firing rate of MD neurons was calculated as the total number of spikes across the full duration of the recording divided by recording duration (time from first to last recorded spike per unit). Similar to other brain regions across species, firing rates of MD units across subdivisions approximate a log-normal distribution (data not shown). Additionally, firing rate was calculated for periods with no bar movement or auditory stimulation by removing spikes 2 seconds before and 5 seconds after a bar push and by removing spikes 100 ms before and 2 seconds after a random tone (mean excluded recording duration: 12.82 ± 0.10 min, mean included recording duration: 21.47 ± 0.14 min). We alternatively calculated firing rate through the median inter-spike interval (ISI) per neuron (1/median ISI). CV was calculated as standard deviation of the ISIs divided by the mean of the ISIs per neuron. CV2 was calculated as 2 x (ISI_n + 1_ – ISI_n_)/ISI_n + 1_ – ISI_n_) per neuron.

### Analysis of burst firing metrics

Burst firing behaviour of MD neurons was calculated as described previously^78^. The threshold for bursting was defined based on the valley between the first two Gaussians of a triple Gaussian mixture model fit of the ISI distribution for all MD units (∼6 ms). Two or more identified spikes with a consecutive ISI lower than 6 ms were classified as bursts. Spikes with ISIs of > 6ms were considered ‘single spikes’. Burst index was calculated by dividing the number of spikes in bursts by the total number of spikes (ratio) across the full recording duration. Burst rate was calculated as the number of bursts across the full recording period (time from first to last recorded spike per unit). Burst duration (ms) was calculated as the time between the first and last spike within a burst. Burst length was calculated as the mean number of spikes per burst. All burst metrics were additionally reanalysed for varying ISI thresholds used to classify bursts (spikes with a consecutive ISI < 4ms and < 10ms).

### Classification

For classification of MD neurons into their anatomical subdivisions, a total of 15 electrophysiological metrics were employed: single-channel waveform features (amplitude, trough-to-peak duration, peak-trough ratio, shoulder asymmetry from refined waveforms), multichannel waveform features (waveform spread, 1/velocity dorsal, 1/velocity ventral), spiking activity (firing rate, firing rate from the ISIs, CV, CV2), and burst firing (burst index, burst rate, burst duration, and burst length). Classification was performed with linear kernel binary support vector machine (SVM) learners. Performance was evaluated using ten-fold cross-validation. For each fold, the classifier was trained on 90% of the data and classifier accuracy was tested on the remaining 10% of the data. Training and test data were z-scored across the entire dataset. To account for unequal numbers of recorded units across MD subdivisions, each class was downsampled to 470 units (determined by the 472 units in MDc that contained valid values across all metrics; MDm: 732 valid units, MDl: 474 valid units) randomly and without replacement. Accuracy was calculated as the proportion of correctly classified neurons across all cross-validation folds and averaged over 100 independent runs of the classifier. Confusion matrices were generated by normalizing by the number of observations in each true class to yield row-wise classification percentages. To assess the relative contribution of the different electrophysiology features to classifier accuracy, we compared the average classifier accuracy restricted to waveform, spiking, or bursting metrics instead. As a control, MD subdivisions labels for each neuron were randomly shuffled across all neurons to estimate chance-level classifier performance.

### Hierarchical clustering

A total of 15 electrophysiological metrics (single-channel waveform features: amplitude, trough-to-peak duration, peak-trough ratio, shoulder asymmetry from refined waveforms, multichannel waveform features: waveform spread, 1/velocity dorsal, 1/velocity ventral, spiking activity: firing rate, firing rate from the ISIs, CV, CV2, and burst firing: burst index, burst rate, burst duration, and burst length) were used to determine whether MD subdivisions segregate into distinct clusters. Hierarchical clustering was performed on z-scored median values for all 1930 MD neurons using 1 – Spearman’s rank correlation coefficient as a distance metric and complete linkage for agglomeration. Optimal leaf ordering was applied to improve dendrogram visualisation. Heatmaps were generated using the z-scored median values for the 15 electrophysiological metrics across all 1930 MD neurons, with rows ordering corresponding to the hierarchical clustering to facilitate visualisation of patterns across MD subdivisions.

The same 15 electrophysiological metrics were used to perform hierarchical clustering of MDm, MDc, MDl, PO, LP, LD, LG, and MGB. Clustering was performed on z-scored median values for the 3843 thalamic units (1930 units from MD) using the same parameters as above. Optimal leaf ordering was applied to improve dendrogram visualisation.

### Statistical analysis

All quantifications and statistical analyses were performed using custom-written code in Matlab v2021b. Where appropriate, data sets were analysed using the Kolmogorov-Smirnov test for normality. Non-normal distributions are accompanied by median values, while normal distributions are accompanied by means. Data sets with normal distributions were analysed for significance using analysis of variance (ANOVA) measures followed by *post hoc* Tukey-Kramer comparisons. Data sets with non-normal distributions were analyzed using Kruskal-Wallis tests followed by *post h*oc Dunn’s test for multiple comparisons. No outlier data were identified or removed unless indicated. Data are expressed as mean/median ±standard error of the mean throughout. Percentage changes are reported relative to the reference subdivision and were calculated as

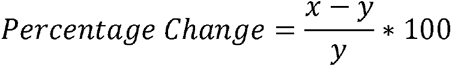

with x denoting the reference subdivision and y denoting the comparison subdivision. Effect size is expressed as η^2^, a measure of the variance in the dataset attributed to differences between the groups. All correlations were examined using Spearman’s rank correlation test. Significance threshold was set at *P* < 0.05. We defined \**P* < 0.05, \*\**P* < 0.01, \*\*\**P* < 0.001. Exact *P*-values values are provided throughout the text, except when *P* < 0.001. Descriptive statistics and statistical test details can be found in **Supplementary Table 2**.

## REFERENCES

1. Mitchell A, Chakraborty S. What does the mediodorsal thalamus do? Front Syst Neurosci. 2013;7:37.

2. Sherman SM. Thalamus plays a central role in ongoing cortical functioning. Nat Neurosci. 2016;19(4):533–41.

3. Bolkan SS, Stujenske JM, Parnaudeau S, Spellman TJ, Rauffenbart C, Abbas AI, et al. Thalamic projections sustain prefrontal activity during working memory maintenance. Nature Neuroscience. 2017;20(7):987–96.

4. Parnaudeau S, O’Neill PK, Bolkan SS, Ward RD, Abbas AI, Roth BL, et al. Inhibition of mediodorsal thalamus disrupts thalamofrontal connectivity and cognition. Neuron. 2013;77(6):1151–62.

5. Schmitt LI, Wimmer RD, Nakajima M, Happ M, Mofakham S, Halassa MM. Thalamic amplification of cortical connectivity sustains attentional control. Nature. 2017;545(7653):219–23.

6. Rikhye RV, Gilra A, Halassa MM. Thalamic regulation of switching between cortical representations enables cognitive flexibility. Nature Neuroscience. 2018;21(12):1753–63.

7. Alcaraz F, Fresno V, Marchand AR, Kremer EJ, Coutureau E, Wolff M. Thalamocortical and corticothalamic pathways differentially contribute to goal-directed behaviors in the rat. eLife. 2018;7:e32517.

8. Parnaudeau S, Taylor K, Bolkan SS, Ward RD, Balsam PD, Kellendonk C. Mediodorsal Thalamus Hypofunction Impairs Flexible Goal-Directed Behavior. Biological psychiatry. 2015;77(5):445–53.

9. Timbie C, Barbas H. Pathways for Emotions: Specializations in the Amygdalar, Mediodorsal Thalamic, and Posterior Orbitofrontal Network. J Neurosci. 2015;35(34):11976–87.

10. Li XB, Inoue T, Nakagawa S, Koyama T. Effect of mediodorsal thalamic nucleus lesion on contextual fear conditioning in rats. Brain research. 2004;1008(2):261–72.

11. The Apulian Network on Risk for Psychosis, Lella A, Antonucci LA, Passiatore R, Bellantuono L, Selvaggi P, et al. Thalamocortical Structural Covariation Networks Are Related to Familial Risk for Schizophrenia in the Context of Lower Nuclei Volume Estimates in Patients: An ENIGMA Study. Biological psychiatry. 2025;98(9):698–711.

12. Stedehouder J, Panti K, Peng Y, Stagg CJ, Sharott A. Higher-Order Thalamus is Pivotal in Schizophrenia-Associated Pathophysiology. bioRxiv. 2026:2026.01.24.699491.

13. Cardoso EF, Maia FM, Fregni F, Myczkowski ML, Melo LM, Sato JR, et al. Depression in Parkinson’s disease: convergence from voxel-based morphometry and functional magnetic resonance imaging in the limbic thalamus. Neuroimage. 2009;47(2):467–72.

14. Li F, Zheng X, Wang H, Meng L, Chen M, Hui Y, et al. Mediodorsal thalamus projection to medial prefrontal cortical mediates social defeat stress-induced depression-like behaviors. Neuropsychopharmacology. 2024;49(8):1318–29.

15. Frau-Méndez MA, Fernández-Vega I, Ansoleaga B, Blanco Tech R, Carmona Tech M, Antonio Del Rio J, et al. Fatal familial insomnia: mitochondrial and protein synthesis machinery decline in the mediodorsal thalamus. Brain Pathol. 2017;27(1):95–106.

16. Wicker E, Hyder SK, Forcelli PA. Pathway-specific inhibition of critical projections from the mediodorsal thalamus to the frontal cortex controls kindled seizures. Prog Neurobiol. 2022;214:102286.

17. Patel S, Millan MH, Meldrum BS. Decrease in excitatory transmission within the lateral habenula and the mediodorsal thalamus protects against limbic seizures in rats. Exp Neurol. 1988;101(1):63–74.

18. Bertram EH. Extratemporal lobe circuits in temporal lobe epilepsy. Epilepsy Behav. 2014;38:13–8.

19. Brumback AC, Ellwood IT, Kjaerby C, Iafrati J, Robinson S, Lee AT, et al. Identifying specific prefrontal neurons that contribute to autism-associated abnormalities in physiology and social behavior. Mol Psychiatry. 2018;23(10):2078–89.

20. Meda KS, Patel T, Braz JM, Malik R, Turner ML, Seifikar H, et al. Microcircuit Mechanisms through which Mediodorsal Thalamic Input to Anterior Cingulate Cortex Exacerbates Pain-Related Aversion. Neuron. 2019;102(5):944–59.e3.

21. Cai S, Huang L, Zou J, Jing L, Zhai B, Ji G, et al. Changes in thalamic connectivity in the early and late stages of amnestic mild cognitive impairment: a resting-state functional magnetic resonance study from ADNI. PLoS One. 2015;10(2):e0115573.

22. Monje MHG, Blesa J, García-Cabezas M, Obeso JA, Cavada C. Changes in thalamic dopamine innervation in a progressive Parkinson’s disease model in monkeys. Mov Disord. 2020;35(3):419–30.

23. Meyer GM, Pines AR, Roldan A, Aracil-Bolaños I, Settle EG, Sahin IA, et al. Deep brain stimulation and psychosis: A case series and two candidate causal brain circuits. medRxiv. 2025.

24. Cascella N, Butala AA, Mills K, Kim MJ, Salimpour Y, Wojtasievicz T, et al. Deep Brain Stimulation of the Substantia Nigra Pars Reticulata for Treatment-Resistant Schizophrenia: A Case Report. Biological psychiatry. 2021;90(10):e57–e9.

25. Zhang DX, Bertram EH. Suppressing limbic seizures by stimulating medial dorsal thalamic nucleus: factors for efficacy. Epilepsia. 2015;56(3):479–88.

26. Mackenzie G, Gilmour W, Yang S-S, Suveges S, MacFarlane J, Kanodia A, et al. Focused ultrasound neuromodulation of mediodorsal thalamus disrupts decision flexibility during reward learning. bioRxiv. 2025:2025.06.03.657634.

27. Krettek JE, Price JL. The cortical projections of the mediodorsal nucleus and adjacent thalamic nuclei in the rat. J Comp Neurol. 1977;171(2):157–91.

28. Groenewegen HJ. Organization of the afferent connections of the mediodorsal thalamic nucleus in the rat, related to the mediodorsal-prefrontal topography. Neuroscience. 1988;24(2):379–431.

29. Mitchell A. The mediodorsal thalamus as a higher order thalamic relay nucleus important for learning and decision-making. Neuroscience & Biobehavioral Reviews. 2015;54:76–88.

30. Klein JC, Rushworth MF, Behrens TE, Mackay CE, de Crespigny AJ, D’Arceuil H, et al. Topography of connections between human prefrontal cortex and mediodorsal thalamus studied with diffusion tractography. Neuroimage. 2010;51(2):555–64.

31. Ordemann GJ, Heckler GJ, Springer K, Driver F, Jackson A, Brumback AC. HCN channel currents underlie distinct neurophysiology of mediodorsal thalamus subnuclei. bioRxiv. 2025:2025.10.05.680564.

32. Alcaraz F, Marchand AR, Courtand G, Coutureau E, Wolff M. Parallel inputs from the mediodorsal thalamus to the prefrontal cortex in the rat. European Journal of Neuroscience. 2016;44(3):1972–86.

33. de Kloet SF, Bruinsma B, Terra H, Heistek TS, Passchier EMJ, van den Berg AR, et al. Bi-directional regulation of cognitive control by distinct prefrontal cortical output neurons to thalamus and striatum. Nature Communications. 2021;12(1):1994.

34. Onishi K, Kikuchi SS, Abe T, Tokuhara T, Shimogori T. Molecular cell identities in the mediodorsal thalamus of infant mice and marmoset. J Comp Neurol. 2022;530(7):963–77.

35. Lyuboslavsky P, Ordemann GJ, Kizimenko A, Brumback AC. Two contrasting mediodorsal thalamic circuits target the mouse medial prefrontal cortex. Journal of neurophysiology. 2024;131(5):876–90.

36. Hoover WB, Vertes RP. Anatomical analysis of afferent projections to the medial prefrontal cortex in the rat. Brain Struct Funct. 2007;212(2):149–79.

37. Kuramoto E, Pan S, Furuta T, Tanaka YR, Iwai H, Yamanaka A, et al. Individual mediodorsal thalamic neurons project to multiple areas of the rat prefrontal cortex: A single neuron-tracing study using virus vectors. Journal of Comparative Neurology. 2017;525(1):166–85.

38. Murphy MJM, Deutch AY. Organization of afferents to the orbitofrontal cortex in the rat. J Comp Neurol. 2018;526(9):1498–526.

39. Ray JP, Price JL. The organization of the thalamocortical connections of the mediodorsal thalamic nucleus in the rat, related to the ventral forebrain-prefrontal cortex topography. J Comp Neurol. 1992;323(2):167–97.

40. Ray JP, Price JL. The organization of projections from the mediodorsal nucleus of the thalamus to orbital and medial prefrontal cortex in macaque monkeys. Journal of Comparative Neurology. 1993;337(1):1–31.

41. Nelson AC, Kapoor V, Vaughn E, Gnanasegaram JA, Rubinstein ND, Talay M, et al. Molecular and neural control of social hierarchy by a forebrain-thalamocortical circuit. Cell. 2025;188(20):5535–54.e23.

42. Phillips JW, Schulmann A, Hara E, Winnubst J, Liu C, Valakh V, et al. A repeated molecular architecture across thalamic pathways. Nature Neuroscience. 2019;22(11):1925–35.

43. Browning PG, Chakraborty S, Mitchell AS. Evidence for mediodorsal thalamus and prefrontal cortex interactions during cognition in macaques. Cerebral cortex. 2015;25(11):4519–34.

44. Chakraborty S, Ouhaz Z, Mason S, Mitchell AS. Macaque parvocellular mediodorsal thalamus: dissociable contributions to learning and adaptive decision-making. Eur J Neurosci. 2019;49(8):1041–54.

45. Jia X, Siegle JH, Bennett C, Gale SD, Denman DJ, Koch C, et al. High-density extracellular probes reveal dendritic backpropagation and facilitate neuron classification. Journal of neurophysiology. 2019;121(5):1831–47.

46. Swadlow HA, Gusev AG. The impact of ‘bursting’ thalamic impulses at a neocortical synapse. Nature Neuroscience. 2001;4(4):402–8.

47. Sherman SM. Tonic and burst firing: dual modes of thalamocortical relay. Trends in neurosciences. 2001;24(2):122–6.

48. Phillips JM, Fish LR, Kambi NA, Redinbaugh MJ, Mohanta S, Kecskemeti SR, et al. Topographic organization of connections between prefrontal cortex and mediodorsal thalamus: Evidence for a general principle of indirect thalamic pathways between directly connected cortical areas. Neuroimage. 2019;189:832–46.

49. Roy DS, Zhang Y, Halassa MM, Feng G. Thalamic subnetworks as units of function. Nature Neuroscience. 2022;25(2):140–53.

50. Li Y, Lopez-Huerta VG, Adiconis X, Levandowski K, Choi S, Simmons SK, et al. Distinct subnetworks of the thalamic reticular nucleus. Nature. 2020;583(7818):819–24.

51. Gao C, Leng Y, Ma J, Rooke V, Rodriguez-Gonzalez S, Ramakrishnan C, et al. Two genetically, anatomically and functionally distinct cell types segregate across anteroposterior axis of paraventricular thalamus. Nature Neuroscience. 2020;23(2):217–28.

52. Oldham S, Ball G. A phylogenetically-conserved axis of thalamocortical connectivity in the human brain. Nature Communications. 2023;14(1):6032.

53. Markram H, Toledo-Rodriguez M, Wang Y, Gupta A, Silberberg G, Wu C. Interneurons of the neocortical inhibitory system. Nature Reviews Neuroscience. 2004;5(10):793–807.

54. Hu H, Gan J, Jonas P. Fast-spiking, parvalbumin+ GABAergic interneurons: From cellular design to microcircuit function. Science. 2014;345(6196):1255263.

55. Pinault D. The thalamic reticular nucleus: structure, function and concept. Brain research reviews. 2004;46(1):1–31.

56. Alloway KD, Hoffer ZS, Hoover JE. Quantitative comparisons of corticothalamic topography within the ventrobasal complex and the posterior nucleus of the rodent thalamus. Brain research. 2003;968(1):54–68.

57. Yamawaki N, Shepherd GM. Synaptic circuit organization of motor corticothalamic neurons. The Journal of Neuroscience. 2015;35(5):2293–307.

58. Perry BAL, Mitchell AS. Considering the Evidence for Anterior and Laterodorsal Thalamic Nuclei as Higher Order Relays to Cortex. Frontiers in Molecular Neuroscience. 2019;12:167.

59. Spring S, Lerch JP, Henkelman RM. Sexual dimorphism revealed in the structure of the mouse brain using three-dimensional magnetic resonance imaging. Neuroimage. 2007;35(4):1424–33.

60. Georgescu IA, Popa D, Zagrean L. The anatomical and functional heterogeneity of the mediodorsal thalamus. Brain Sciences. 2020;10(9):624.

61. Akam T, Lustig A, Rowland JM, Kapanaiah SKT, Esteve-Agraz J, Panniello M, et al. Open-source, Python-based, hardware and software for controlling behavioural neuroscience experiments. eLife. 2022;11:e67846.

62. Wang Q, Ding S-L, Li Y, Royall J, Feng D, Lesnar P, et al. The Allen Mouse Brain Common Coordinate Framework: A 3D Reference Atlas. Cell. 2020;181(4):936–53.e20.

63. Birman D, Yang KJ, West SJ, Karsh B, Browning Y, Laboratory tIB, et al. Pinpoint: trajectory planning for multi-probe electrophysiology and injections in an interactive web-based 3D environment. bioRxiv. 2023:2023.07.14.548952.

64. Siegle JH, López AC, Patel YA, Abramov K, Ohayon S, Voigts J. Open Ephys: an open-source, plugin-based platform for multichannel electrophysiology. J Neural Eng. 2017;14(4):045003.

65. Shamash P, Carandini M, Harris K, Steinmetz N. A tool for analyzing electrode tracks from slice histology. bioRxiv. 2018:447995.

66. Petersen PC, Siegle JH, Steinmetz NA, Mahallati S, Buzsáki G. CellExplorer: A framework for visualizing and characterizing single neurons. Neuron. 2021;109(22):3594–608.e2.

67. Jun JJ, Steinmetz NA, Siegle JH, Denman DJ, Bauza M, Barbarits B, et al. Fully integrated silicon probes for high-density recording of neural activity. Nature. 2017;551(7679):232–6.

68. Chen S, Liu Y, Wang ZA, Colonell J, Liu LD, Hou H, et al. Brain-wide neural activity underlying memory-guided movement. Cell. 2024;187(3):676–91.e16.

69. Liu LD, Chen S, Hou H, West SJ, Faulkner M, Economo MN, et al. Accurate Localization of Linear Probe Electrode Arrays across Multiple Brains. eNeuro. 2021;8(6).

70. Peters AJ, Fabre JMJ, Steinmetz NA, Harris KD, Carandini M. Striatal activity topographically reflects cortical activity. Nature. 2021;591(7850):420–5.

71. Paxinos G, Franklin KB. Paxinos and Franklin’s the mouse brain in stereotaxic coordinates: Academic press; 2019.

72. Groenewegen H, Witter M. Thalamus. The Rat Nervous System. 2004:407–53.

73. Pachitariu M, Steinmetz N, Kadir S, Carandini M, Harris K. Fast and accurate spike sorting of high-channel count probes with KiloSort. Proceedings of the 30th International Conference on Neural Information Processing Systems; Barcelona, Spain: Curran Associates Inc.; 2016. p. 4455–63.

74. Steinmetz NA, Zatka-Haas P, Carandini M, Harris KD. Distributed coding of choice, action and engagement across the mouse brain. Nature. 2019;576(7786):266–73.

75. Julie M.J. Fabre EHvB, Andrew J. Peters, Matteo Carandini, & Kenneth D. Harris Bombcell: automated curation and cell classification of spike-sorted electrophysiology data (1.0.0). Zenodo 2023.

76. Lopes-Dos-Santos V, Brizee D, Dupret D. Spatio-temporal organization of network activity patterns in the hippocampus. Cell Rep. 2025;44(6):115808.

77. Buzsaki G, Kandel A. Somadendritic backpropagation of action potentials in cortical pyramidal cells of the awake rat. Journal of neurophysiology. 1998;79(3):1587–91.

78. Mizuseki K, Royer S, Diba K, Buzsáki G. Activity dynamics and behavioral correlates of CA3 and CA1 hippocampal pyramidal neurons. Hippocampus. 2012;22(8):1659–80.

