## Supplementary Materials for "Distinct In Vivo Electrophysiological Profiles of Mediodorsal Thalamus Subdivisions"

Katerina Panti *et al*.

**The PDF file includes:**

Figures S1 to S10
Tables S1 and S3

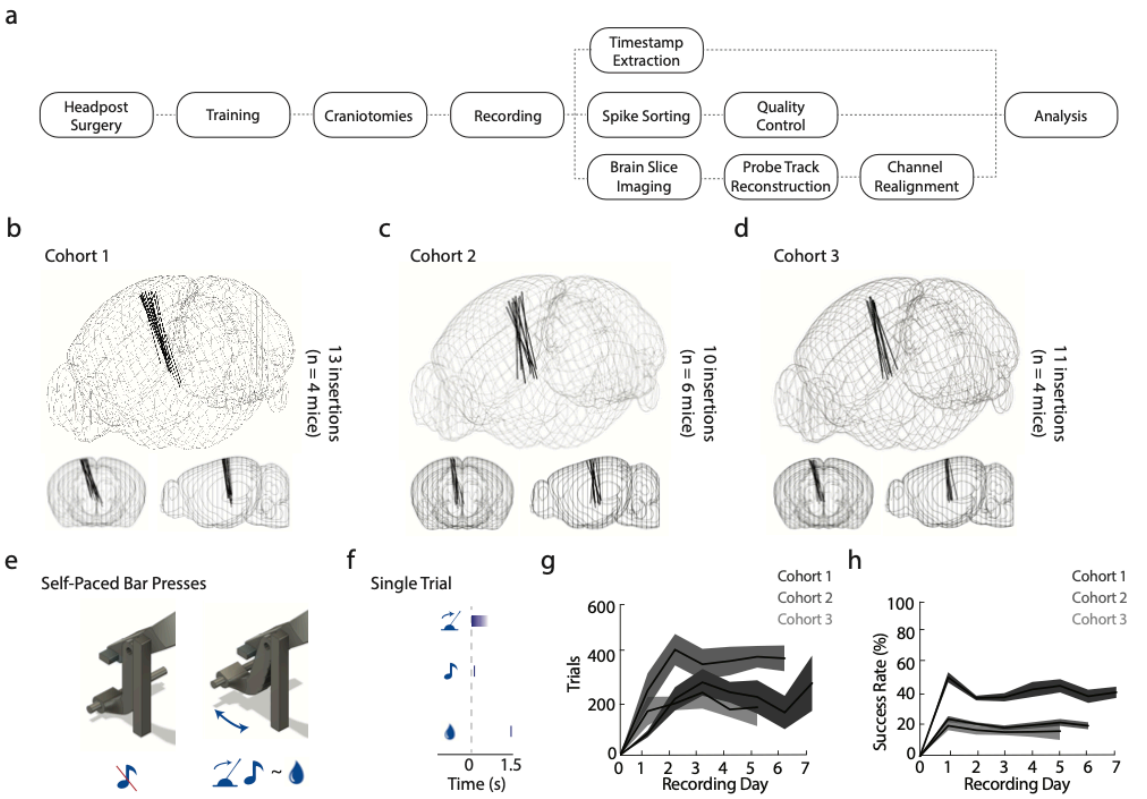


**Supplementary Figure 1. *In vivo* electrophysiological recordings across three wildtype cohorts and under a sensorimotor behavioural task**

**(a)** Experimental pipeline. Each cohort of animals underwent surgery, habituation, training, and recording, followed by a standardized reconstruction and analysis pipeline.

**(b)** 3D schematic of brains showing reconstructed probe trajectories (black lines) for cohort 1 (*n* = 4 mice).

**(c)** 3D schematic of brains showing reconstructed probe trajectories (black lines) for cohort 2 (*n* = 6 mice).

**(d)** 3D schematic of brains showing reconstructed probe trajectories (black lines) for cohort 3 (*n* = 4 mice).

**(e, f)** Behavioral task setup. Adult, male, head-fixed mice were trained to press a bar self-paced for delayed reward. Bar press onset was accompanied by an auditory cue (100 ms, 2-16 kHz, tone frequency was set separately for each animal).

**(g)** Number of trials across recording days for the three cohorts in different shades of grey.

**(h)** Success rate across recording days for the three cohorts in different shades of grey (percentage of trials where the animal pushed the metal bar and returned to the home position within 250-1000ms). Shaded areas denote standard error of the means.

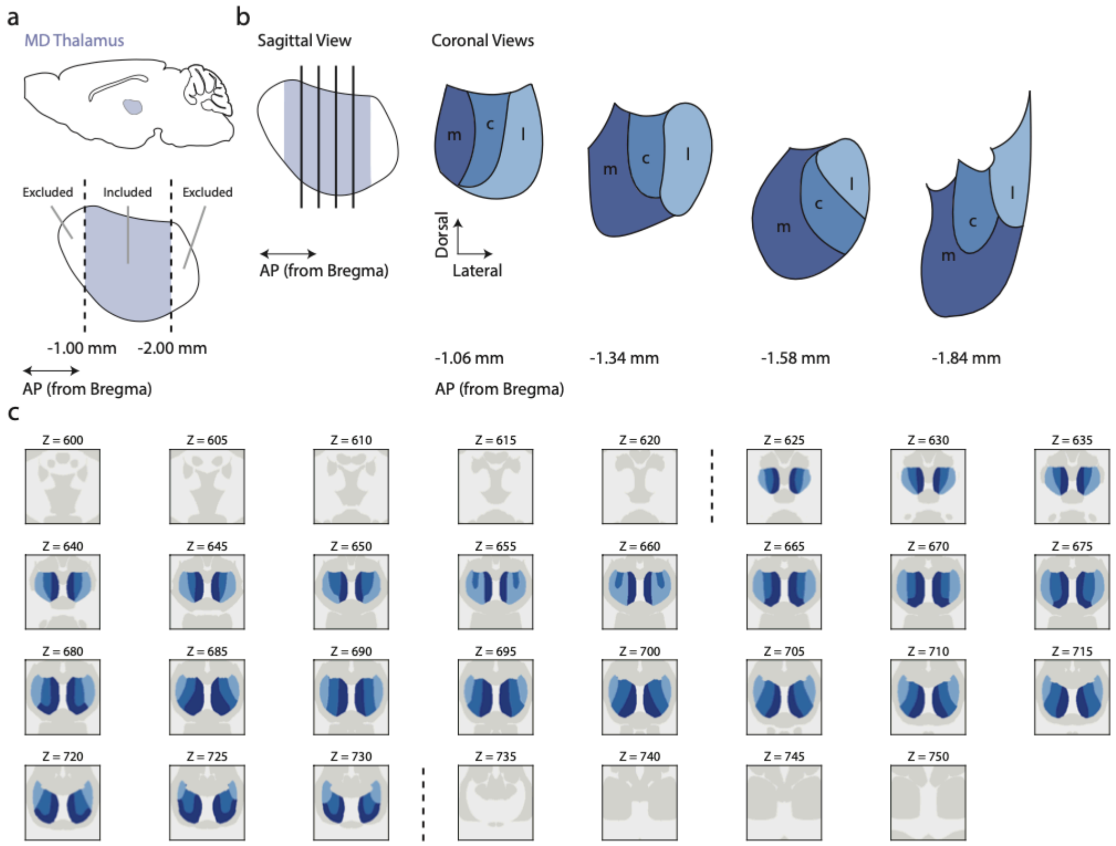


**Supplementary Figure 2. Atlas of MD subdivisions**

**(a)** Sagittal schematic of location of rodent MD thalamus (*top*, blue) and anteroposterior (AP) axis borders (*bottom*) included in subsequent analysis. Most anterior and posterior portion of MD thalamus were not included in analysis.

**(b)** Sagittal overview similar to (a) and coronal schematics of rodent MD across four locations across the AP axis in right hemisphere. MDm ('m', dark blue); MDc ('c', blue); MDl ('l', light blue).

**(c)** Coronal view of the mouse atlas using the Allen CCF (Shamash et al), adapted to classify MDm (dark blue), MDc (blue), and MDl (light blue) across the AP axis of MD. Numbers indicate CCF AP coordinates.

AP antero-posterior.

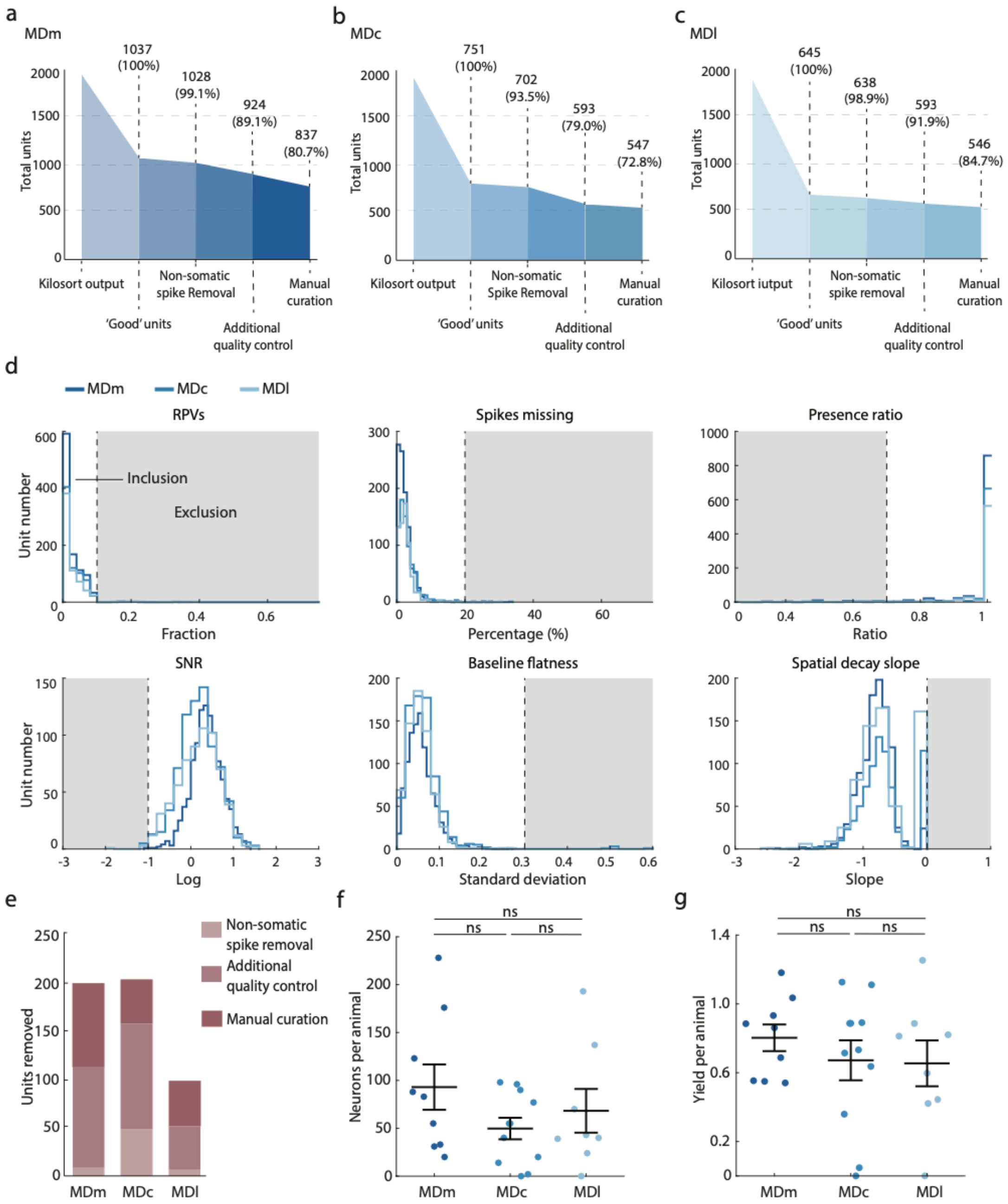


**Supplementary Figure 3. Quality control for mediodorsal thalamus units**

**(a)** Flowchart showing quality control steps and neuron exclusion based on successive criteria for neurons in MDm.

**(b)** Same as (b) for MDc.

**(c)** Same as (b) for MDl.

**(d)** Histograms of quality control characteristics in before quality control ($n$ = 2433 neurons from 14 mice) along with exclusion criteria (vertical black dashed lines) and to be further excluded neurons (grey shaded areas).

**(e)** Number of neurons removed for various stages of quality control for each MD subdivision.

**(f)** Comparison of recorded number of neurons per animal across MDm (*n* = 9), MDc (*n* = 11), and MDl (*n* = 8). No difference occurred in recorded number of neurons per subdivision per animal.

(g) Comparison of yield (number neurons per channel) per animal across MDm (*n* = 9), MDc (*n* = 11), and MDl (*n* = 8). No difference occurred in yield per subdivision per animal.

Black lines in (f) and (g) indicate means. Error bars denote s.e.m.

ns $P > 0.05$. One-way ANOVA.

Statistics in **Supplementary Table 2**.

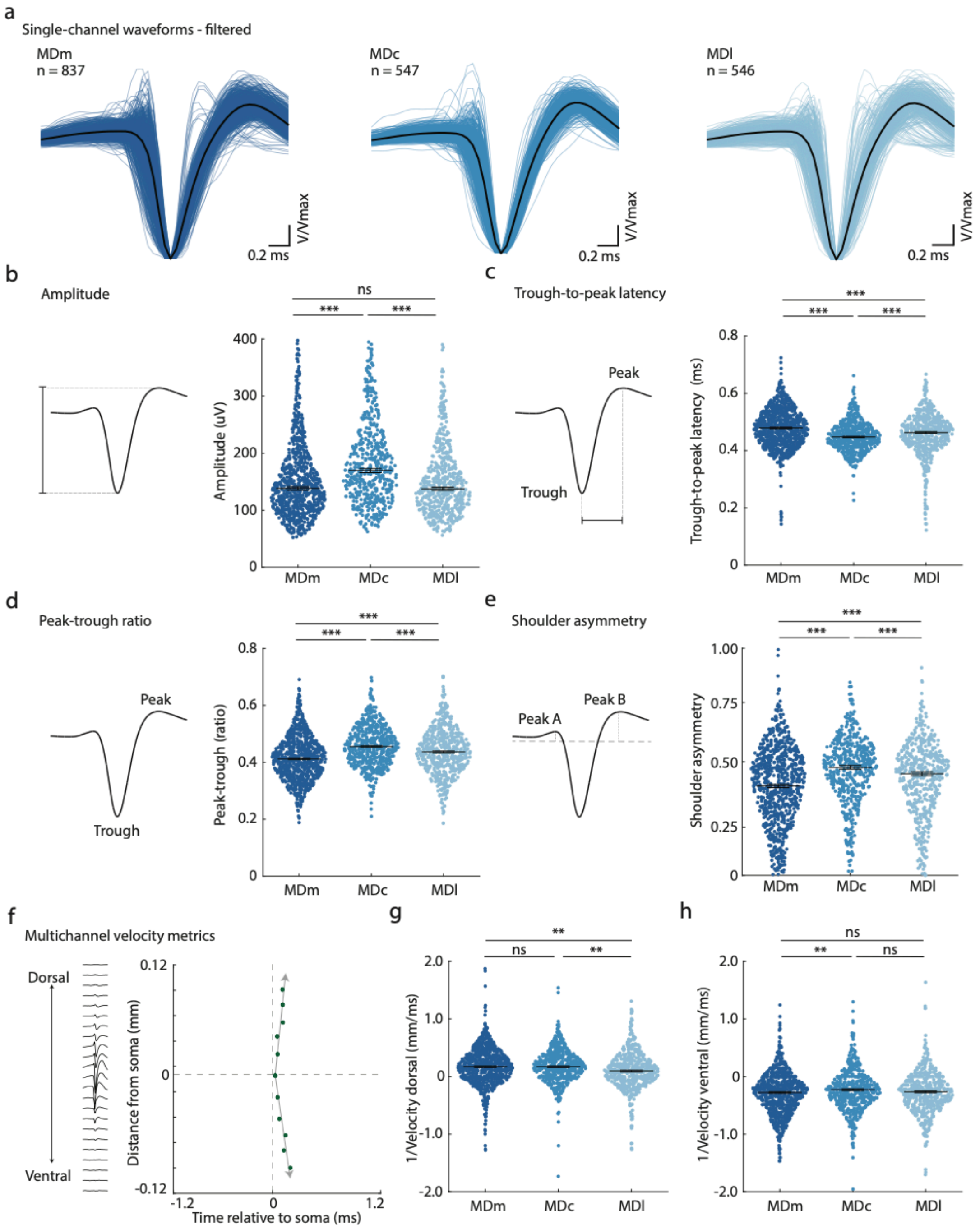


**Supplementary Figure 4. Mediodorsal thalamus subdivision differences in waveform metrics persist in high-pass filtered waveforms**

**(a)** Single-channel high-pass filtered waveforms normalized to their peak amplitude of individual neurons in MDm (dark blue), MDc (blue), and MDl (light blue). Black line indicates mean of all single-channel filtered waveforms.

**(b)** *Left:* Schematic of extracellular waveform amplitude calculation. *Right*: Swarm plot comparing amplitude across MD subdivisions, showing differences across subdivisions. Y-axis capped at 400 µV for clarity.

**(c)** *Left:* Schematic of trough-to-peak latency calculation. *Right*: Swarm plot comparing trough-to-peak latency across MD subdivisions, showing differences across subdivisions.

**(d)** *Left:* Schematic of peak-trough ratio calculation. *Right*: Swarm plot comparing peak-trough ratio across MD subdivisions, showing differences across subdivisions.

**(e)** *Left:* Schematic of shoulder asymmetry calculation. *Right*: Swarm plot comparing shoulder asymmetry across MD subdivisions, showing differences across subdivisions.

**(f)** Multichannel waveform along the probe for example neuron (left) and propagation trajectory along the probe (right). For each electrode location (y axis, y = 0 indicates soma location) the time of the waveform trough relative to the trough recorded from the soma is plotted on the x-axis. Dorsal velocity (above 0) and ventral velocity (below 0) are calculated separately using linear regression (grey arrows).

**(g)** Swarm plot comparing the inverse of dorsal velocity across MD subdivisions, showing minor differences across subdivisions.

**(h)** Swarm plot comparing the inverse of ventral velocity across MD subdivisions, showing minor differences across subdivisions.

Dots represent individual neurons. Black lines in (b) through (e) indicate medians. Black lines in (g) and (h) indicate means. Error bars denote s.e.m.

n.s. $P > 0.05$, * $P < 0.05$, ** $P < 0.01$, *** $P < 0.001$. In (b-e) Kruskal-Wallis test followed by Dunn's multiple comparisons. In (g, h): One-way ANOVA test followed by *post hoc* Tukey-Kramer comparisons.

Statistics in **Supplementary Table 2**.

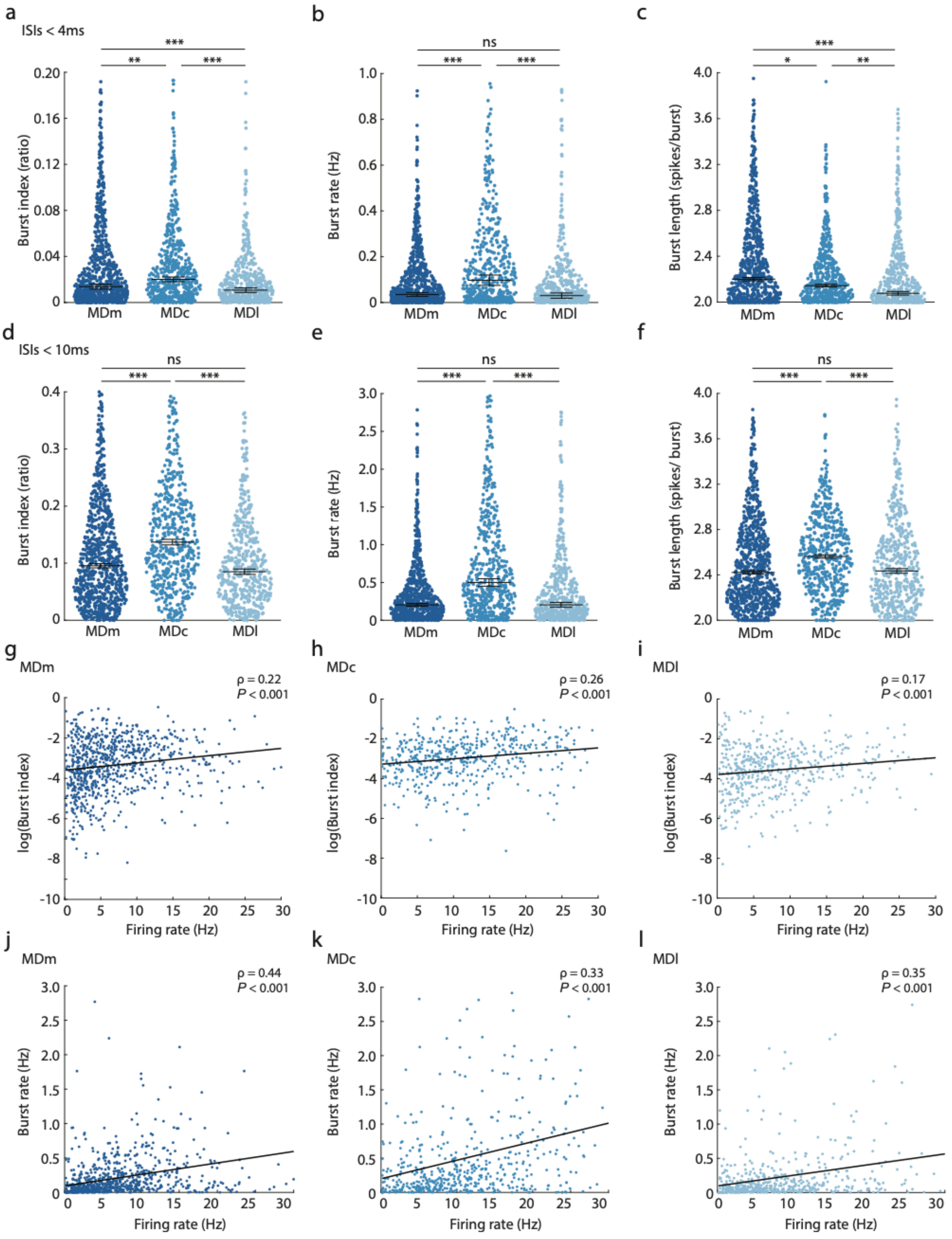


**Supplementary Figure 5. Differences in burst firing metrics across MD subdivisions with varying inter-spike interval thresholds**

**(a)** Swarm plot comparing burst length across MD subdivisions for ISIs < 4ms, showing differences across subdivisions. MDm: $n$ = 796, MDc: $n$ = 531, MDl: $n$ = 519.

**(b)** Swarm plot comparing burst index across MD subdivisions for ISIs < 4ms, showing differences across subdivisions.

**(c)** Swarm plot comparing burst rate across MD subdivisions for ISIs < 4ms, showing differences across subdivisions.

**(d)** Same as (a) for ISIs < 10ms. MDm: $n = 832$, MDc: $n = 541$, MDl: $n = 541$, showing differences across subdivisions.

**(e)** Same as (b) for ISIs < 10ms.

**(f)** Same as (c) for ISIs < 10ms.

**(g)** Scatter plot depicting log burst index as a function of firing rate for MDm cells, supporting a positive relation between spiking and bursting.

**(h)** Same as (g) for MDc cells.

**(i)** Same as (g) for MDl cells.

**(j)** Scatter plot depicting burst rate as a function of firing rate for MDm cells, supporting a positive relation between spiking and bursting.

**(k)** Same as (j) for MDc cells.

**(l)** Same as (j) for MDl cells.

Blue dots represent individual neurons. Black lines in (a-f) indicate medians. Error bars denote s.e.m. Black lines in (g-i) indicate linear regression.

n.s. $P > 0.05$, * $P < 0.05$, ** $P < 0.01$, *** $P < 0.001$. In (a-f) Kruskal-Wallis test followed by Dunn's multiple comparisons. In (g-l) Spearman's correlation (ρ).

Statistics in **Supplementary Table 2**

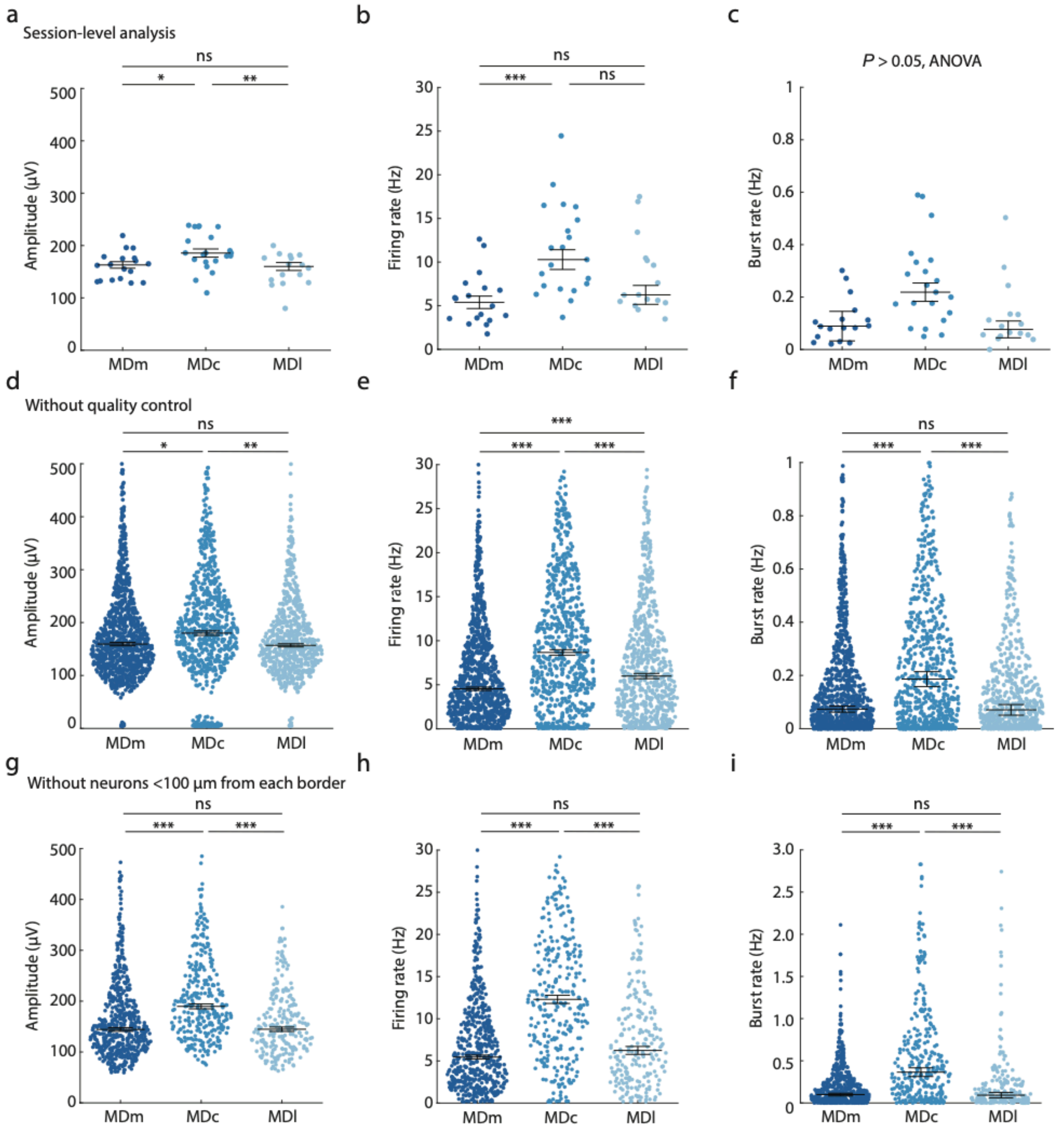


**Supplementary Figure 6. Robust differences in electrophysiological metrics across MD subdivisions**

**(a)** Swarm plot comparing amplitude with neurons averaged for each recording session across MD subdivisions. Each dot represents median amplitude for one session. MDm: $n$ = 18, MDc: $n$ = 21, MDl: $n$ = 16.

**(b)** Swarm plot comparing firing rate for each session across MD subdivisions.

**(c)** Swarm plot comparing burst rate for each session across MD subdivisions.

**(d)** Swarm plot comparing amplitude across MD subdivisions prior to quality control and manual curation. MDm: $n$ = 1037, MDc: $n$= 751, MDl: $n$ = 645.

**(e)** Swarm plot comparing firing rate across MD subdivisions prior to quality control and manual curation.

**(f)** Swarm plot comparing burst rate across MD subdivisions prior to quality control and manual curation.

**(g)** Swarm plot comparing amplitude across MD subdivisions for units >100 µm from each anatomical border between MD subdivisions and with other brain regions. MDm: $n$ = 563, MDc: $n$ = 305, MDl: $n$ = 225.

**(h)** Swarm plot comparing firing rate across MD subdivisions for units >100 µm from each anatomical border between MD subdivisions and with other brain regions.

**(i)** Swarm plot comparing burst rate across MD subdivisions for units >100 µm from each anatomical border between MD subdivisions and with other brain regions.

Blue dots represent session averages in (a) through (c) and individual neurons in (d) through (i). Black lines in (a-c) indicate means. Black lines in (d-i) indicate medians. Error bars denote s.e.m.

*** $P < 0.001$; ** $P < 0.01$; ns non-significant $P > 0.05$. In (a) through (c): One-way ANOVA followed by *post hoc* Tukey-Kramer comparisons. In (d) through (i): Kruskal-Wallis test followed by Dunn's multiple comparisons.

Statistics in **Supplementary Table 2**.

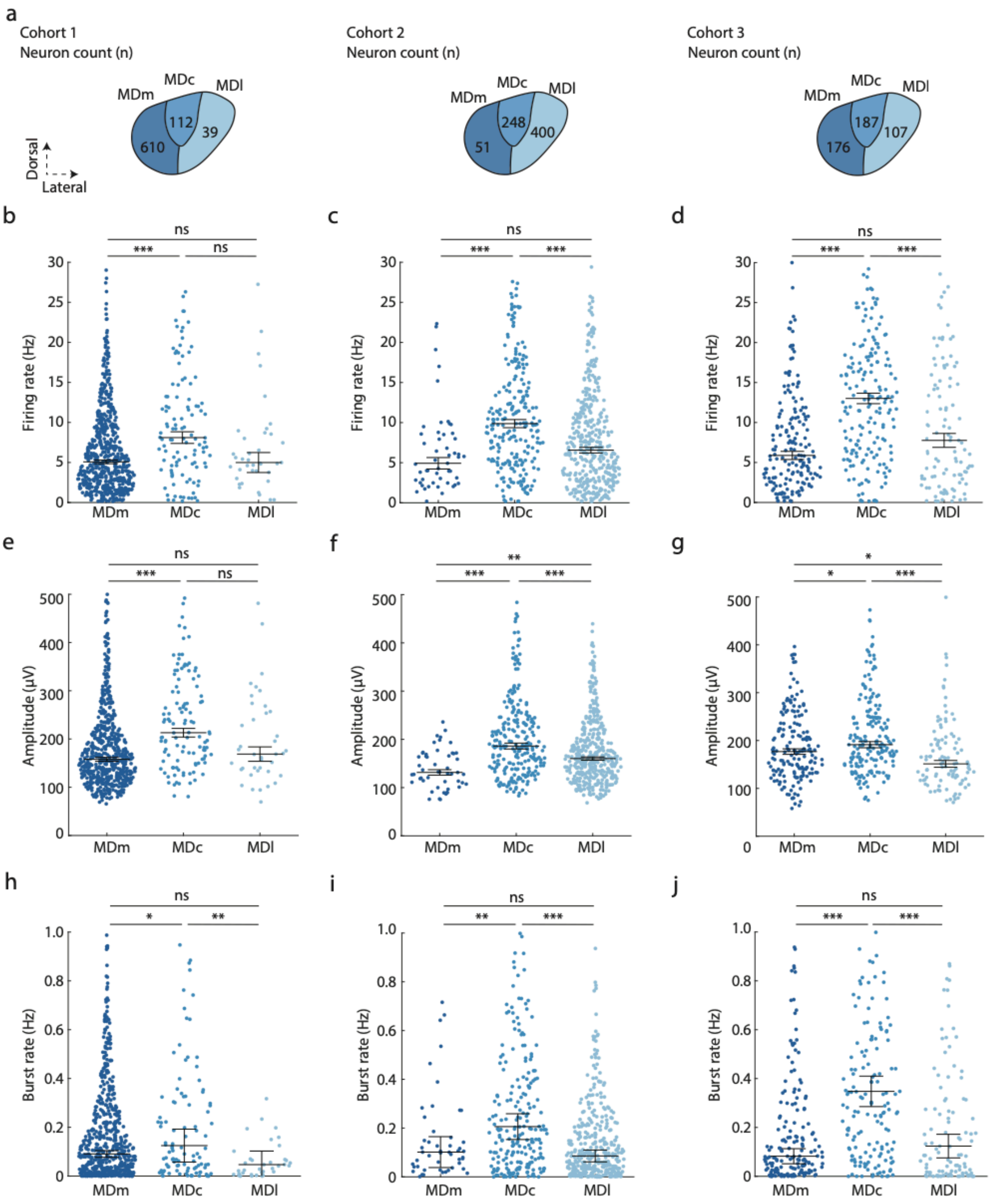


**Supplementary Figure 7. Differences in electrophysiological metrics of MD subdivisions across the three independent cohorts**

**(a)** Neuron count for each MD subdivision across the three cohorts.

**(b)** Swarm plot comparing firing rate across MD subdivisions for cohort 1.

**(c)** Same as (a) for cohort 2.

**(d)** Same as (a) for cohort 3.

**(e)** Swarm plot comparing amplitude across MD subdivisions for cohort 2.

**(f)** Same as (d) for cohort 2.

**(g)** Same as (d) for cohort 3.

**(h)** Swarm plot comparing burst rate across MD subdivisions for cohort 3.

**(i)** Same as (g) for cohort 2.

**(j)** Same as (g) for cohort 3.

Blue dots represent individual neurons. Black lines in (b) through (j) indicate medians. Error bars denote standard error of the mean.

*** $P < 0.001$; ** $P < 0.01$; * $P < 0.05$; ns non-significant $P > 0.05$. Kruskal-Wallis test followed by Dunn's multiple comparisons.

Statistics in **Supplementary Table 2**.

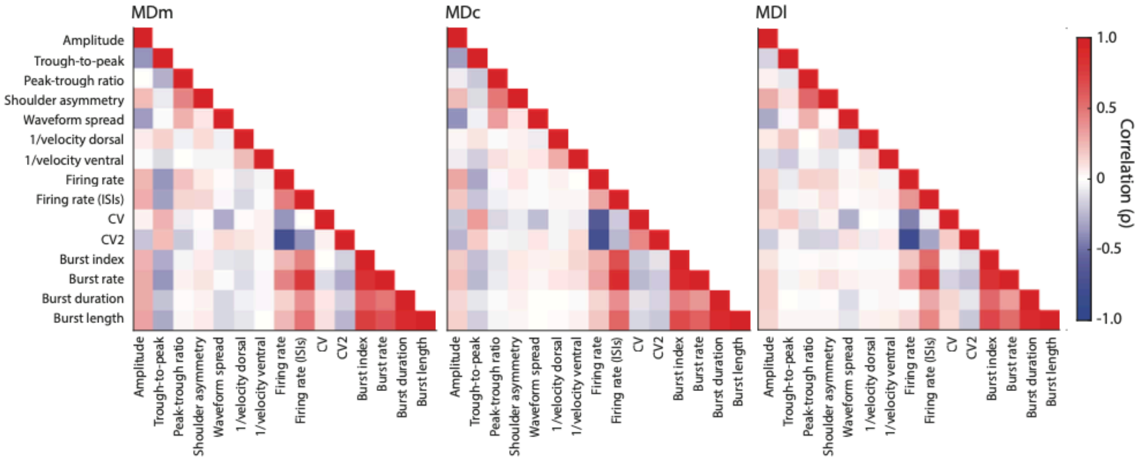


**Supplementary Figure 8. Cross-correlation of electrophysiological metrics**

Spearman correlation heatmap for 15 normalized electrophysiology metrics in MDm, MDc, and MDl. Single-channel waveform metrics include amplitude, trough-to-peak latency, peak-trough ratio, shoulder asymmetry. Multichannel waveform metrics include waveform spread, 1/velocity dorsal, 1/velocity ventral. Spiking metrics include firing rate, firing rate calculated from the ISIs (1/median ISI per unit), CV, CV2. Burst firing metrics include burst index, burst rate, burst duration, burst length.

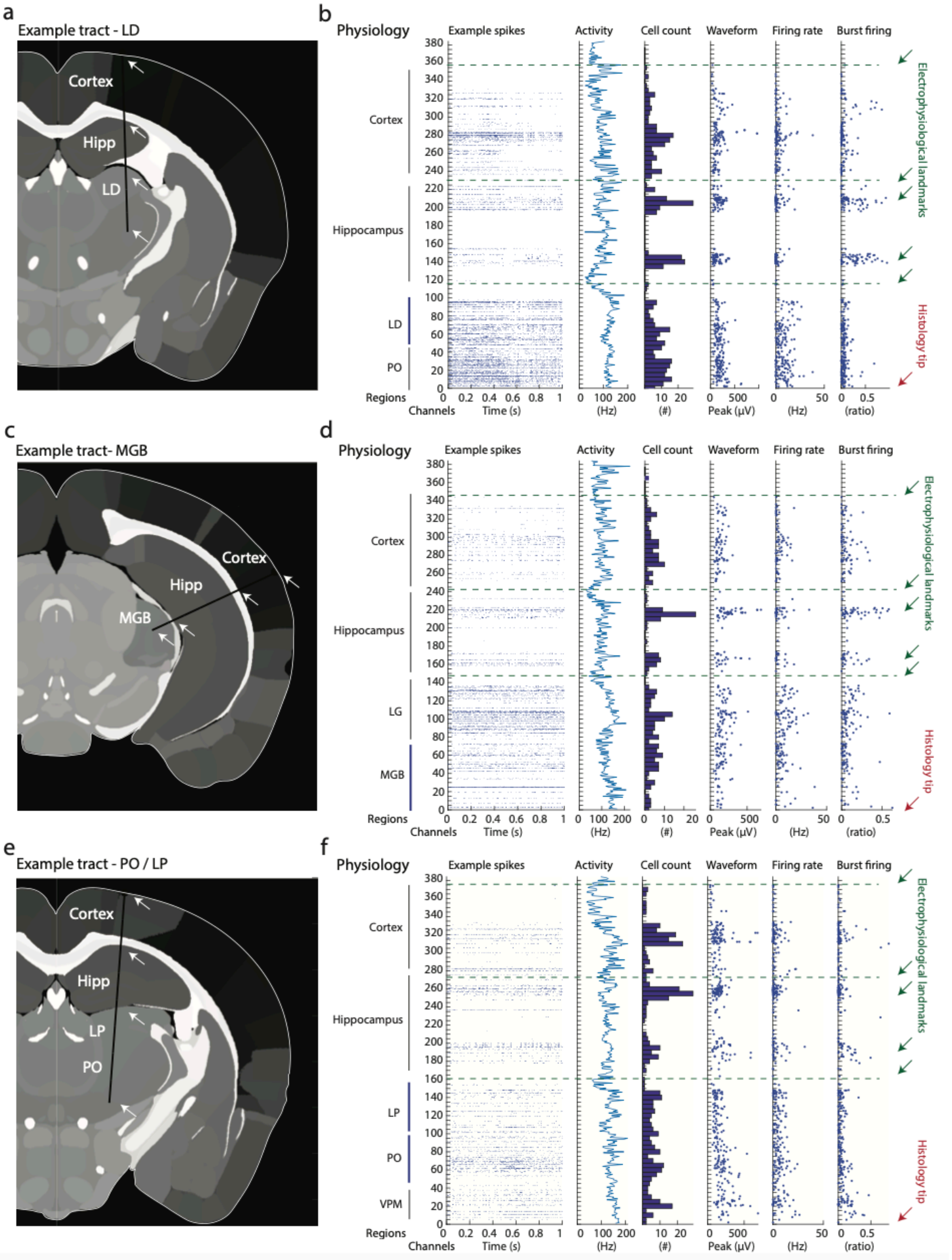


**Supplementary Figure 9. Probe channel realignment for additional non-MD thalamic nuclei**

**(a)** Example coronal view of an insertion track of a Neuropixels probe (black line) to LD thalamus with traversed regions indicated in text and landmarks indicated with white arrows.

**(b)** Example region alignment for the single probe insertion shown in (a). Information on spiking, waveform, and bursting activity across channel depth through cortex, hippocampus, LD, and PO.

**(c)** Example coronal view of an insertion track of a Neuropixels probe (black line) to MGB thalamus with traversed regions indicated in text and landmarks indicated with white arrows. Note that the probe was inserted through left hemisphere, and this image is flipped for visualization purposes.

**(d)** Example region alignment for the single probe insertion shown in (c). Information on spiking, waveform, and bursting activity across channel depth through cortex, hippocampus, LG, and MGB.

**(e)** Example coronal view of an insertion track of a Neuropixels probe (black line) to PO/LP thalamus with traversed regions indicated in text and landmarks indicated with white arrows.

**(f)** Example region alignment for the single probe insertion shown in (e). Information on spiking, waveform, and bursting activity across channel depth through cortex, hippocampus, LP, PO, and VPM.

In (b), (d), (f): Red lines indicate landmarks for rearranging channels for brain region borders. Green arrows represent previously validated electrophysiological landmarks used to rearrange channels.

LD laterodorsal nucleus, LG lateral geniculate nucleus, LP lateroposterior nucleus, MGB medial geniculate body, PO posterior nucleus, VPM ventral posterior nucleus of the thalamus.

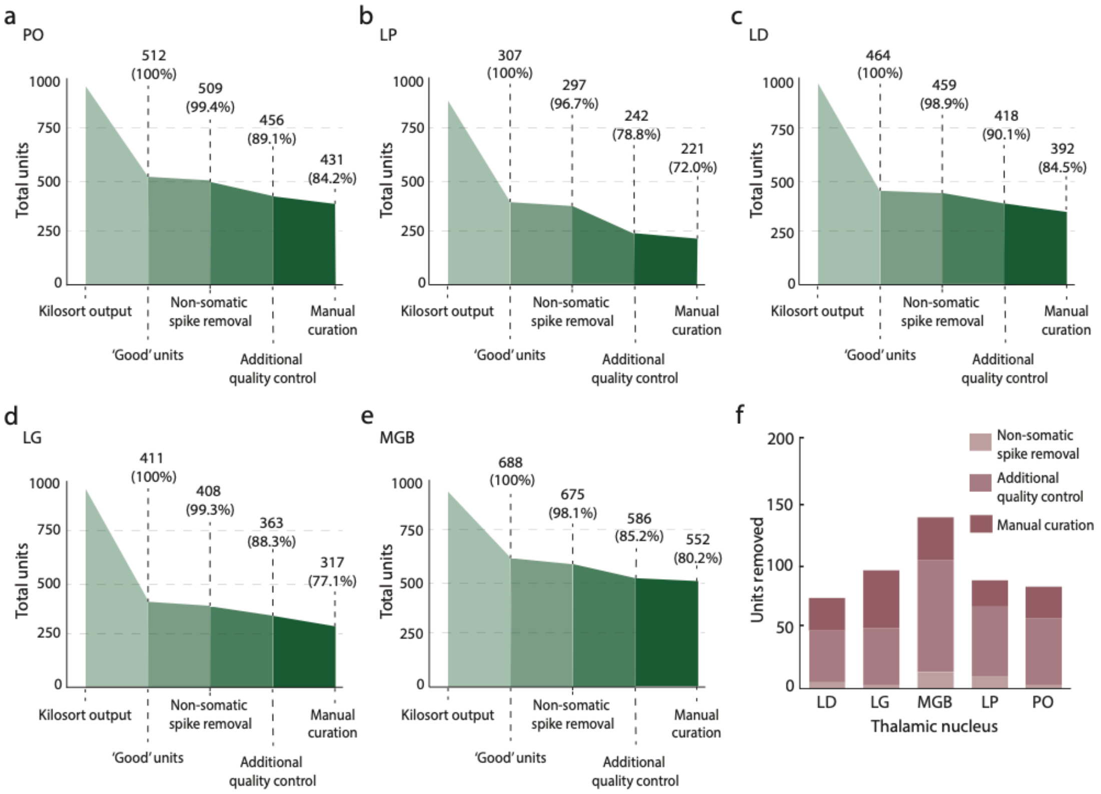


**Supplementary Figure 10. Quality control for non-MD thalamic nuclei**

**(a)** Flowchart showing quality control steps and neuron exclusion based on successive criteria for neurons in PO.

**(b)** Same as (a) for LP.

**(c)** Same as (a) for LD.

**(d)** Same as (a) for LG.

**(e)** Same as (a) for MGB.

**(f)** Number of neurons removed for various stages of quality control for each thalamic nucleus.

LD laterodorsal nucleus, LG lateral geniculate nucleus, LP lateroposterior nucleus, MGB medial geniculate body, PO posterior nucleus.

**Supplementary Table 1. Number of recorded units per MD subdivision across sessions and animals**

Number of units per MD subdivision recorded for each session across animals (*n* = 12) and cohorts (*n* = 3). No units retained from animal 5 and animal 8 following automated and manual curation.

| Cohort | Animal | Recording session | MDm units | MDc units | MDl units |
|---|---|---|---|---|---|
| 1 | 1 | 1 | 87 | 18 | 0 |
| 1 | 1 | 2 | 36 | 2 | 0 |
| 1 | 2 | 1 | 12 | 0 | 0 |
| 1 | 2 | 2 | 1 | 42 | 10 |
| 1 | 2 | 3 | 32 | 48 | 29 |
| 1 | 2 | 4 | 38 | 0 | 0 |
| 1 | 3 | 1 | 24 | 0 | 0 |
| 1 | 3 | 2 | 131 | 0 | 0 |
| 1 | 3 | 3 | 73 | 0 | 0 |
| 1 | 4 | 1 | 0 | 2 | 0 |
| 1 | 4 | 2 | 39 | 0 | 0 |
| 1 | 4 | 3 | 79 | 0 | 0 |
| 1 | 4 | 4 | 58 | 0 | 0 |
| 2 | 6 | 1 | 0 | 46 | 21 |
| 2 | 6 | 2 | 0 | 5 | 101 |
| 2 | 6 | 3 | 0 | 4 | 71 |
| 2 | 7 | 1 | 0 | 0 | 46 |
| 2 | 7 | 2 | 31 | 89 | 0 |
| 2 | 7 | 3 | 0 | 9 | 24 |
| 2 | 9 | 1 | 20 | 55 | 0 |
| 3 | 10 | 1 | 0 | 6 | 5 |
| 3 | 10 | 2 | 0 | 14 | 57 |
| 3 | 10 | 3 | 0 | 20 | 75 |
| 3 | 11 | 1 | 0 | 0 | 43 |
| 3 | 12 | 1 | 0 | 0 | 22 |
| 3 | 12 | 2 | 55 | 6 | 0 |
| 3 | 12 | 3 | 0 | 43 | 17 |
| 3 | 12 | 4 | 0 | 28 | 1 |
| 3 | 13 | 1 | 0 | 5 | 12 |
| 3 | 13 | 2 | 0 | 9 | 12 |
| 3 | 13 | 3 | 33 | 0 | 0 |
| 3 | 14 | 1 | 0 | 32 | 0 |
| 3 | 14 | 2 | 78 | 0 | 0 |
| 3 | 14 | 3 | 10 | 64 | 0 |

**Supplementary Table 2. Statistics**

Statistical overview. Included as a separate file.

**Supplementary Table 3. Number of recorded units per thalamic nucleus across sessions and animals**

Number of units recorded from PO, LP, LD, LG, and MGB for each session across animals ($n$ = 14) and cohorts ($n$ = 3).

LD laterodorsal nucleus, LG lateral geniculate nucleus, LP lateroposterior nucleus, MGB medial geniculate body, PO posterior nucleus.

| Cohort | Animal | Recording session | PO units | LP units | LD units | LG units | MGB units |
|---|---|---|---|---|---|---|---|
| 1 | 1 | 1 | 0 | 1 | 0 | 0 | 0 |
| 1 | 1 | 2 | 0 | 13 | 0 | 6 | 0 |
| 1 | 1 | 3 | 0 | 14 | 0 | 0 | 0 |
| 1 | 2 | 1 | 0 | 9 | 0 | 0 | 0 |
| 1 | 2 | 2 | 0 | 3 | 0 | 0 | 0 |
| 1 | 2 | 3 | 0 | 10 | 0 | 0 | 0 |
| 1 | 2 | 4 | 0 | 2 | 0 | 0 | 0 |
| 1 | 3 | 1 | 0 | 21 | 0 | 0 | 0 |
| 1 | 3 | 2 | 0 | 0 | 0 | 47 | 3 |
| 1 | 3 | 3 | 0 | 0 | 0 | 38 | 1 |
| 1 | 3 | 4 | 0 | 13 | 0 | 2 | 0 |
| 1 | 4 | 1 | 0 | 5 | 0 | 35 | 0 |
| 1 | 4 | 2 | 0 | 8 | 0 | 0 | 0 |
| 1 | 4 | 3 | 0 | 8 | 0 | 15 | 0 |
| 1 | 4 | 4 | 0 | 2 | 0 | 0 | 0 |
| 1 | 4 | 5 | 0 | 16 | 0 | 0 | 0 |
| 2 | 5 | 1 | 0 | 0 | 0 | 46 | 16 |
| 2 | 5 | 2 | 70 | 0 | 67 | 0 | 0 |
| 2 | 5 | 3 | 0 | 2 | 0 | 27 | 5 |
| 2 | 5 | 4 | 147 | 0 | 71 | 0 | 0 |
| 2 | 5 | 5 | 0 | 0 | 0 | 62 | 49 |
| 2 | 5 | 6 | 80 | 0 | 35 | 0 | 0 |
| 2 | 5 | 7 | 0 | 0 | 0 | 18 | 0 |
| 2 | 5 | 8 | 0 | 0 | 0 | 0 | 39 |
| 2 | 6 | 1 | 0 | 0 | 0 | 0 | 27 |
| 2 | 6 | 2 | 3 | 15 | 0 | 0 | 0 |
| 2 | 6 | 3 | 0 | 5 | 0 | 0 | 0 |
| 2 | 7 | 1 | 0 | 0 | 0 | 0 | 15 |
| 2 | 7 | 2 | 0 | 21 | 0 | 0 | 0 |
| 2 | 7 | 3 | 0 | 0 | 0 | 0 | 15 |
| 2 | 7 | 4 | 0 | 0 | 3 | 0 | 0 |
| 2 | 7 | 5 | 122 | 45 | 0 | 0 | 0 |
| 2 | 7 | 6 | 0 | 0 | 0 | 0 | 18 |
| 2 | 7 | 7 | 0 | 6 | 0 | 0 | 0 |

| | | | | | | | |
|---|---|---|---|---|---|---|---|
| 2 | 7 | 8 | 0 | 0 | 0 | 0 | 21 |
| 2 | 8 | 1 | 0 | 0 | 21 | 0 | 0 |
| 2 | 8 | 2 | 0 | 0 | 7 | 0 | 0 |
| 2 | 9 | 1 | 0 | 0 | 57 | 0 | 0 |
| 2 | 9 | 2 | 9 | 0 | 43 | 0 | 0 |
| 3 | 10 | 1 | 0 | 0 | 0 | 16 | 4 |
| 3 | 10 | 2 | 0 | 0 | 0 | 1 | 35 |
| 3 | 10 | 3 | 0 | 0 | 0 | 4 | 3 |
| 3 | 11 | 1 | 0 | 0 | 14 | 0 | 0 |
| 3 | 11 | 2 | 0 | 0 | 0 | 0 | 29 |
| 3 | 11 | 3 | 0 | 0 | 18 | 0 | 0 |
| 3 | 11 | 4 | 0 | 0 | 0 | 0 | 11 |
| 3 | 11 | 5 | 0 | 0 | 4 | 0 | 0 |
| 3 | 11 | 6 | 0 | 0 | 0 | 0 | 11 |
| 3 | 11 | 7 | 0 | 0 | 21 | 0 | 0 |
| 3 | 11 | 8 | 0 | 0 | 0 | 0 | 14 |
| 3 | 11 | 9 | 0 | 0 | 2 | 0 | 0 |
| 3 | 12 | 1 | 0 | 0 | 0 | 0 | 5 |
| 3 | 13 | 1 | 0 | 2 | 0 | 0 | 0 |
| 3 | 13 | 2 | 0 | 0 | 0 | 0 | 22 |
| 3 | 13 | 3 | 0 | 0 | 10 | 0 | 0 |
| 3 | 14 | 1 | 0 | 0 | 0 | 0 | 68 |
| 3 | 14 | 2 | 0 | 0 | 0 | 0 | 42 |
| 3 | 14 | 3 | 0 | 0 | 0 | 0 | 65 |
| 3 | 14 | 4 | 0 | 0 | 19 | 0 | 0 |
| 3 | 14 | 5 | 0 | 0 | 0 | 0 | 34 |
